# MCU gain-and loss-of-function models define the duality of mitochondrial calcium uptake in heart failure

**DOI:** 10.1101/2023.04.17.537222

**Authors:** Joanne F. Garbincius, Timothy S. Luongo, Jonathan P. Lambert, Adam S. Mangold, Emma K. Murray, Alycia N. Hildebrand, Pooja Jadiya, John W. Elrod

## Abstract

**Background:** Mitochondrial calcium (_m_Ca^2+^) uptake through the mitochondrial calcium uniporter channel (mtCU) stimulates metabolism to meet acute increases in cardiac energy demand. However, excessive _m_Ca^2+^ uptake during stress, as in ischemia-reperfusion, initiates permeability transition and cell death. Despite these often-reported acute physiological and pathological effects, a major unresolved controversy is whether mtCU-dependent _m_Ca^2+^ uptake and long-term elevation of cardiomyocyte _m_Ca^2+^ contributes to the heart’s adaptation during sustained increases in workload.

**Objective:** We tested the hypothesis that mtCU-dependent _m_Ca^2+^ uptake contributes to cardiac adaptation and ventricular remodeling during sustained catecholaminergic stress.

**Methods:** Mice with tamoxifen-inducible, cardiomyocyte-specific gain (αMHC-MCM x flox-stop-MCU; MCU-Tg) or loss (αMHC-MCM x *Mcu*^fl/fl^; *Mcu*-cKO) of mtCU function received 2-wk catecholamine infusion.

**Results:** Cardiac contractility increased after 2d of isoproterenol in control, but not *Mcu*-cKO mice. Contractility declined and cardiac hypertrophy increased after 1-2-wk of isoproterenol in MCU-Tg mice. MCU-Tg cardiomyocytes displayed increased sensitivity to Ca^2+^- and isoproterenol-induced necrosis. However, loss of the mitochondrial permeability transition pore (mPTP) regulator cyclophilin D failed to attenuate contractile dysfunction and hypertrophic remodeling, and increased isoproterenol-induced cardiomyocyte death in MCU-Tg mice.

**Conclusions:** mtCU _m_Ca^2+^ uptake is required for early contractile responses to adrenergic signaling, even those occurring over several days. Under sustained adrenergic load excessive MCU-dependent _m_Ca^2+^ uptake drives cardiomyocyte dropout, perhaps independent of classical mitochondrial permeability transition pore opening, and compromises contractile function. These findings suggest divergent consequences for acute versus sustained _m_Ca^2+^ loading, and support distinct functional roles for the mPTP in settings of acute _m_Ca^2+^ overload versus persistent _m_Ca^2+^ stress.

## INTRODUCTION

Rapid uptake of calcium (Ca^2+^) into the mitochondrial matrix occurs via the mitochondrial calcium uniporter channel (mtCU), a multi-protein complex that spans the inner mitochondrial membrane^1, 2^ and consists of a pore-forming subunit, MCU^3, 4^, and the accessory regulatory proteins EMRE^5–7^; MCUB^8^, MICU1 and MICU2/3^9–15^, and MCUR1^16, 17^. Cardiomyocyte mitochondrial Ca^2+^ (_m_Ca^2+^) uptake through the mtCU increases when the local cytosolic Ca^2+^ concentration rises past ∼400nM^1^. The mtCU is thus responsible for rapidly increasing net _m_Ca^2+^ content in response to acute elevations in cytosolic Ca^2+^ concentration, such as occurs when the heart is subjected to acute sympathetic stimulation and cellular Ca^2+^ cycling is enhanced. This acute increase in _m_Ca^2+^ content is thought to act as a second messenger that stimulates TCA cycle dehydrogenases and a subsequent increase in the rate of mitochondrial ATP production. This process guarantees that ATP generation is increased in parallel with the stimulation of ATP-consuming cytosolic processes such as myofilament crossbridge cycling that drive the contraction of the heart^2^.

Recent investigations indicate that physiological _m_Ca^2+^ uptake through the mtCU is required to support an increased heart rate and increased cardiac contractility in the minutes immediately following acute β-adrenergic stimulation^18–20^. However, _m_Ca^2+^ uptake through the mtCU also contributes to _m_Ca^2+^ overload and mitochondrial permeability transition (mPT) in response to extreme cytosolic Ca^2+^ levels^21^. The observation that conditional deletion of *Mcu* in adult mouse cardiomyocytes is sufficient to prevent _m_Ca^2+^ overload and subsequent necrotic cell death during cardiac ischemia-reperfusion injury^19, 20^ supports this model. Yet, despite such evidence for the contribution of mtCU-dependent _m_Ca^2+^ uptake to the heart’s response to acute physiological or pathological Ca^2+^ stress, the relevance of the mtCU and sustained elevations in _m_Ca^2+^ concentration to the heart’s responses to chronic increases in cardiac workload remains controversial.

Cardiac MCU expression is increased in patients with pressure overload due to aortic stenosis and in mouse models of cardiac hypertrophy ^22, 23^. The expression of additional components of the _m_Ca^2+^ exchange machinery, including the mtCU regulator MICU1 and the mitochondrial sodium-calcium exchanger NCLX, are also altered in the failing human heart^24, 25^, suggestive of altered _m_Ca^2+^ handling in chronic heart disease. Such findings have prompted investigation into whether mtCU-dependent _m_Ca^2+^ uptake may be an adaptive or maladaptive aspect of the heart’s response to chronic stress. Early experiments in mice showed no positive nor detrimental effect of constitutive, global *Mcu* deletion or adult, cardiomyocyte-specific *Mcu* deletion on functional decompensation or pathological cardiac remodeling following chronic pressure overload (transverse aortic constriction; TAC)^20, 26^. More recent conflicting studies indicate that pharmacologic blockade of MCU protects against declines in cardiac function after TAC^23^; or instead that increasing _m_Ca^2+^ uptake in pressure-+ neurohormonal overload-induced heart failure via viral MCU overexpression can be beneficial by reducing oxidative stress and ultimately improving contractile function^27^. Direct comparison of these disparate results, and understanding of the ultimate functional consequences of sustained uniporter-dependent _m_Ca^2+^ uptake in chronic heart disease, is confounded by differences in experimental approaches used to manipulate mtCU function (pharmacologic vs. genetic; global vs. cell-type specific); the time available for compensatory adaptations to occur in models with constitutive vs. inducible gene disruption; distinct timepoints of experimental interventions throughout the course of cardiac decompensation (prior to onset of precipitating insult vs. after the appearance of contractile dysfunction); model system (species; assessment of cardiomyocyte vs. whole-heart function); and inclusion of single vs. multiple analysis timepoints among these studies^28^. Therefore, we performed longitudinal studies comparing the impact of inducible, adult cardiomyocyte-specific loss-or gain-of uniporter function to clarify the role that the mtCU and altered _m_Ca^2+^ content play in the heart’s adaptation and eventual maladaptation to chronic catecholamine stimulation and elevated cardiac workload.

Here, we demonstrate that the mtCU is required to increase cardiac contractility over the first several days of isoproterenol stimulation and conclude that any mtCU-independent _m_Ca^2+^ uptake that may occur over this time frame is insufficient to support *in vivo* increases in cardiac workrate. We find that increased _m_Ca^2+^ uptake through the mtCU becomes detrimental only after 1-2 wks of sustained catecholaminergic stimulation, suggesting that persistent _m_Ca^2+^ loading results in a gradual shift from the _m_Ca^2+^ signal mediating beneficial early metabolic adaptation to an increased cardiac workload, to the _m_Ca^2+^ signal later driving maladaptive responses. We further show that genetic disruption of mitochondrial permeability transition is insufficient to prevent the functional decline or pathological remodeling of MCU-overexpressing hearts subjected to chronic adrenergic stress, and increases isoproterenol-induced cardiomyocyte death. Our findings suggest that although _m_Ca^2+^ uptake through the mtCU is required to support initial increases in contractility at the onset of adrenergic stimulation, under prolonged Ca^2+^ stress, _m_Ca^2+^ uptake drives cardiomyocyte dropout, possibly independent of classical cyclophilin D-regulated cell death, and compromises cardiac function.

## METHODS

### Mice

To generate a conditional MCU overexpression mouse model, the coding sequence and stop codon of human *MCU* cDNA (NCBI reference sequence NM_138357.1) was cloned into a custom CAG-loxP-CAT-loxP plasmid following a strategy we described previously^29^. The resulting construct contained the artificial CAG promoter followed by loxP sites flanking chloramphenicol acetyltransferase (CAT) with multiple stop sequences, with the *MCU* cDNA following the second loxP site. Cre-mediated recombination of the loxP sites excises the CAT-stop sequence, allowing for the strong ubiquitous CAG promoter to drive expression of the MCU transgene. The plasmid was linearized and injected into 1-cell embryos, which were transplanted into pseudo-pregnant female mice. Resulting flox-stop-MCU founders were crossed to αMHC-Cre mice (Jackson Laboratories strain #009074)^30^ to enable constitutive expression in cardiomyocytes. Founder lines were evaluated for expression and leakiness of the MCU transgene via Western blot. A flox-stop-MCU founder line exhibiting strong, Cre-dependent MCU transgene expression in the heart, and no expression in the absence of Cre, was selected for further use (**Supplemental Fig. S1A**). For tamoxifen inducible, cardiomyocyte-specific transgene expression, flox-stop-MCU mice were crossed to α-myosin heavy chain-MerCreMer mice (αMHC-MCM; “MCM”; Jackson Laboratories strain #005657) to generate αMHC-MCM x flox-stop-MCU (“MCU-Tg”) animals. *Mcu*^fl/fl^ mice were crossed to αMHC-MCM mice to generate αMHC-MCM x Mcu*^fl/fl^*(*Mcu*-cKO) animals and allow tamoxifen-inducible, cardiomyocyte-specific *Mcu* deletion as described and validated previously^19^. The generation of mice with global deletion of cyclophilin D (*Ppif*^-/-^) has been described elsewhere^31^. *Ppif*^-/-^ mice were crossed to MCM mice and to MCU-Tg mice. *Ppif*^+/-^ offspring were interbred to generate MCM and MCU-Tg mice on both wild-type and *Ppif*^-/-^ backgrounds. All mouse lines were maintained on a C57BL/6N background (Jackson Laboratories strain #005304).

### Mice were genotyped for presence of the MCU transgene using one of two primer sets

The first primer set, with expected product size of 599 base pairs, consisted of the forward primer: 5’- CAGTTCACACTCAAGCCTATCT-3’ and the reverse primer: 5’- CTGTCTCTGGCTTCTGGATAAA-3’. The second primer set, with expected product size of 354 base pairs, consisted of the forward primer: 5’- CTGTTGTGCCCTCTGATGAT-3’ and the reverse primer: 5’- GTTGCTGGACCAATGTCTTTAC-3’. PCR reaction mixture contained 1uL tail DNA in DirectPCR Lysis reagent (Viagen Biotech #102-T), 1x Taq buffer (Syd Labs #MB042-EUT), 80µM each dNTPs (New England Biolabs #N0447L), 800nM each forward and reverse primers, and 1.25 U Taq polymerase (Syd Labs #MB042-EUT). The PCR conditions were denaturation at 95°C for 3 minutes, followed by 40 cycles (95°C for 30 seconds, 61°C for 30 seconds, 72°C for 30 seconds), followed by 5 minutes at 72°C.

Mice were genotyped for the presence of the αMHC-Cre or the αMHC-MCM transgene using the forward primer: 5’-GGCGTTTTCTGAGCATACCT-3’ and the reverse primer: 5’- CTACACCAGAGACGGAAATCCA-3’, with an expected product size of 565-585 base pairs.

PCR reaction mixture contained 1uL tail DNA in DirectPCR Lysis reagent, 1x Taq buffer, 80µM each dNTPs, 800nM each forward and reverse primers, and 1.25 U Taq polymerase. The PCR conditions were, denaturation at 95°C for 3 minutes, followed by 34 cycles (95°C for 30 seconds, 55°C for 30 seconds, 72°C for 30 seconds), followed by 10 minutes at 72°C.

Mice were genotyped for *Mcu* using the forward primer: 5’- GAAGGCCTCCTGTTATGGAT-3’ and the reverse primer: 5’-CCAGCTTGGTGAAGCCTGAT-3’, with expected product sizes of 261 base pairs for the wild-type allele and 354 base pairs for the floxed allele. PCR reaction mixture contained 1uL tail DNA in DirectPCR Lysis reagent, 1x Taq buffer, 100µM each dNTPs, 100mM betaine (Sigma-Aldrich #B0300), 1µM each forward and reverse primers, and 1.25 U Taq polymerase. The PCR conditions were: denaturation at 95°C for 3 minutes, followed by 34 cycles (95°C for 30 seconds, 55°C for 30 seconds, 72°C for 30 seconds), followed by 10 minutes at 72°C.

Mice were genotyped for *Ppif* using the forward primers: 5’- CTCTTCTGGGCAAGAATTGC-3’ (wild-type allele) or 5’-GGCTGCTAAAGCGCATGCTCC-3’ (null allele); and the reverse primer: 5’-ATTGTGGTTGGTGAAGTCGCC-3’, with expected product sizes of 850 base pairs for the wild-type allele and 600 base pairs for the null allele. PCR reaction mixture contained 1uL tail DNA in DirectPCR Lysis reagent, 1x Taq buffer, 200µM each dNTPs, 1µM each forward and reverse primers, and 2.5 U Taq polymerase. The PCR conditions were: denaturation at 95°C for 3 minutes, followed by 34 cycles (95°C for 30 seconds, 56°C for 1 minute, 72°C for 1.5 minutes), followed by 5 minutes at 72°C.

All mice were used between 8-25 weeks of age. Both male and female mice were included. Experiments were performed and analyzed using a numbered ear-tagging system to blind the experimenter to mouse genotype and experimental group. All animal experiments followed AAALAC guidelines and were approved by Temple University’s IACUC.

### Tamoxifen-inducible MCU overexpression or *Mcu* deletion

For initial characterization of temporal overexpression of MCU, MCU-Tg mice received intraperitoneal injections with 20mg/kg/day of tamoxifen (Sigma-Aldrich #T5648) dissolved in corn oil (Sigma-Aldrich # C8267) for 5 consecutive days. Control αMHC-MCM mice were subjected to this same tamoxifen injection paradigm. Mice were allowed 2-3 days of tamoxifen washout prior to acute *in vitro* studies. To compare loss-and gain-of MCU function *in vivo*, αMHC-MCM, *Mcu*-cKO, and MCU-Tg mice all received intraperitoneal injections with 40mg/kg/day of tamoxifen dissolved in corn oil for 5 consecutive days to ensure effective deletion of *Mcu*^19^. For *in vivo* experiments, mice were allowed 16 days of tamoxifen washout following the last tamoxifen dose, prior to baseline echocardiography and osmotic minipump implantation.

### Isolation of adult mouse cardiomyocytes

Cardiomyocytes were isolated from the ventricles of adult mouse hearts based on methods previously reported^19, 32^. In brief, mice were injected with 1000U heparin (McKesson Corporation, #691115) and the hearts rapidly excised. The aorta was canulated and the heart perfused with perfusion buffer (120.4mM NaCl (Sigma-Aldrich #S9888), 14.7mM KCl (Amresco #0395), 0.6mM KH_2_PO_4_ (Amresco #0781), 0.6mM NaH_2_PO_4_ (Amresco #0404), 1.2mM MgSO_4_·7H_2_O (Sigma-Aldrich # 230391), 10mM HEPES (Research Products International #H75030), 4.6mM NaHCO_3_ (Sigma-Aldrich #S5761 check), 5.5mM glucose (Sigma-Aldrich #G8270), 10mM 2,3-butanedione monoxime (Sigma-Aldrich #B0753), and 30mM taurine (Sigma-Aldrich #T8691), pH = 7.4), then digested with perfusion buffer supplemented with 1mg/mL collagenase B (Sigma-Aldrich #11088831001), 139µg/mL trypsin (ThermoFisher #15090046) and 12.5µM CaCl_2_ (Sigma-Aldrich # 21115). Digested heart tissue was teased into small pieces using fine forceps and further dissociated by gentle pipetting to release cardiomyocytes. Trypsin digestion was terminated by transferring cells to stopping buffer (perfusion buffer supplemented with 10% fetal bovine serum (Peak Serum #PS-FB3) and 12.5µM CaCl_2_). All cardiomyocytes were used within 3 hours of isolation.

### Mitochondrial isolation

Mitochondria were isolated from hearts of adult mice 1wk after the start of tamoxifen injections based on the approach of Frezza et al.^33^. Excised hearts were minced in ice-cold 1x phosphate buffered saline (PBS) (Morganville Scientific # PH0200) supplemented with 10mM EDTA (BioWORLD #40520000-1), washed 3 times, and digested on ice in 1x PBS supplemented with 10mM EDTA and 83.3 µg/mL trypsin. Digested tissue was rinsed 3 times in 1x PBS supplemented with 10mM EDTA then centrifuged at 200g for 5min at 4°C to pellet the tissue. Tissue chunks were resuspended in ice-cold IBM1 buffer (67mM sucrose (BioWORLD # 41900152-2), 5mM Tris/HCl (BioPioneer #C0116), 5mM KCl, 1mM EDTA, 0.2% bovine serum albumin (Sigma-Aldrich # A3803), pH = 7.2) and homogenized using a glass/Teflon homogenizer with an overhead stirrer (Heidolph Instruments #501-64010-00) at 2000rpm. The homogenate was centrifuged at 700g for 10min at 4°C, and the supernatant centrifuged again at 7200g for 12min at 4°C to pellet mitochondria. Mitochondria were washed in ice-cold IBM2 buffer (250mM sucrose, 0.3mM EGTA/Tris (Sigma-Aldrich #E3889; Amresco #0497); 1mM Tris/HCl, pH = 7.2) and centrifuged at 7200g for 12min at 4°C. The supernatant was removed and isolated mitochondria were used for size exclusion chromatography or for mitochondrial swelling assays as described below.

### Fast protein size-exclusion liquid chromatography (FPLC)

For each replicate experiment, cardiac mitochondria isolated from 3-5 pooled hearts of each genotype were lysed on ice for 30 min in 1X RIPA buffer (EMD Millipore #20-188) supplemented with 1X protease inhibitors (Sigma-Aldrich #S8830-20TAB), and lysates were cleared by centrifuging at 14000g for 10min at 4°C. Protein concentration was determined by bicinchoninic acid assay (BioWORLD #20831001). 2500µg of cleared mitochondrial lysate were fractionated by gel filtration using fast protein size-exclusion liquid chromatography (AKTA Pure FPLC; GE Healthcare), using a Superdex 200 Increase 10/300 column (Sigma-Aldrich, #GE28-9909-44) equilibrated in 1X PBS, at a flow rate of 0.5mL/min. 0.5mL protein fractions were collected, concentrated to 75µL with 3kD molecular weight cutoff AMICON Ultra-0.5 centrifugal filter devices (EMD Millipore #UFC500396) following the manufacturer’s instructions. Concentrated protein fractions were used for western blotting under reducing conditions as described below. Molecular weights of FPLC fractions were calibrated using gel filtration markers (Sigma-Aldrich #MWGF1000).

### Western blotting

Isolated adult mouse cardiomyocytes were pelleted by centrifugation at 200g for 5min, then washed in 1X PBS, and centrifuged again at 200g for 5min. The supernatant was removed, and cardiomyocyte pellets were snap frozen in liquid nitrogen until use. Cardiomyocyte pellets were lysed in ice-cold 1X RIPA buffer supplemented with 1X protease inhibitors and 1X Phosstop phosphatase inhibitor (Roche #04906837001) and sonicated for 10sec. Lysates were centrifuged for 5min at 5000g at 4°C, and the supernatant was used for western blotting. Mouse heart tissue was homogenized in ice-cold 1X RIPA buffer supplemented with 1X protease inhibitors and 1X Phosstop phosphatase inhibitor using a bead mill homogenizer (VWR, #75840–022). Homogenates were sonicated for 10sec, then centrifuged for 5min at 5000g at 4°C, and the supernatant was used for western blotting. Protein concentration in cardiomyocyte or heart lysates was determined by bicinchoninic acid assay. 5X SDS sample buffer (250 µM Tris/HCl, pH 7.0; 40% (v/v) glycerol (Sigma-Aldrich #G5516); 8% (w/v) sodium dodecyl sulfate (Amresco #0227); 20% (v/v) β-mercaptoethanol (Sigma-Aldrich # M6250); 0.1% (w/v) bromophenol blue (Fisher Scientific #BP115-25) was added to samples to a final concentration of 1X, and 20-50µg of protein/well was separated on 10% (w/v) polyacrylamide Tris-glycine SDS gels. For samples from isoproterenol infusion cohorts, 25µg protein/well was separated on NuPAGE 4-12% Bis-Tris midi protein gels (ThermoFisher #WG1403BOX). 20µl of concentrated protein fractions from FPLC experiments were mixed with 5X SDS sample to a final concentration of 1X, and equal volumes of each fraction were separated on NuPAGE 4-12% Bis-Tris midi protein gels.

After separation by electrophoresis, proteins were transferred to polyvinylidene fluoride membranes (Millipore #IPFL00010). Membranes were blocked in blocking buffer (Rockland #MB-070) for 1 hour at room temperature, and then incubated overnight at 4°C in primary antibodies diluted in 50% blocking buffer / 50% Tris buffered saline (bioWORLD #42020056–3) + 0.1% TWEEN (Sigma-Aldrich #P9416) (TBS-T). Primary antibodies and dilutions included: rabbit monoclonal against MCU (Cell Signaling Technologies #14997) 1:1000; rabbit polyclonal against MCU (Sigma #HPA016480) 1:1000; mouse monoclonal against total OXPHOS complexes (Abcam #ab110413) 1:1000; rabbit polyclonal against phospho-PDH E_1_a-Ser^293^ (Abcam #ab92696) 1:1000; mouse monoclonal against PDH E_1_a subunit (Abcam #ab110330) 1:1000; mouse monoclonal against ATP5A (Abcam #ab14748), 1:2000; rabbit polyclonal against MICU1 (Sigma #HPA037480), 1:1000; rabbit polyclonal against EMRE (Bethyl Laboratories # A300-BL19208) 1:1000; mouse monoclonal against cyclophilin D (Abcam #ab110324) 1:1,000; and mouse monoclonal against total PDH subunits (Abcam #ab110416) 1:500. Membranes were washed 3 times in TBS-T and incubated for 1.5 hours at room temperature in secondary antibodies diluted 1:10,000 in 50% blocking buffer / 50% TBS-T. Secondary antibodies and dilutions included: IRDye 800CW goat anti-rabbit (LI-COR #925-32211) 1:10,000; IRDye 800CW Goat anti-Mouse (LI-COR, #926-32210) 1:10,000; IRDye 680RD goat anti-rabbit (LI-COR #926-68071) 1:10,000; and IRDye 680RD goat anti-mouse (LI-COR #925-68070) 1:10,000. Membranes were then washed 3 times in TBS-T and imaged using a LI-COR Odyssey infrared imaging system. All full-length western blots are shown in Supplemental Figs. S1-S5. Densitometric quantification of western blots was performed with LI-COR Image Studio software (LI-COR, version 2.0.38).

### _m_Ca^2+^ handling assays

Adult mouse cardiomyocytes were isolated as described above and counted in stopping buffer before assessment of _m_Ca^2+^ uptake and mitochondrial calcium retention capacity based on methods described previously^14, 19^. For each assay replicate, 300,000 live cardiomyocytes were pelleted at 100g for 3 min, resuspended in extracellular-like Ca^2+^-free buffer (120mM NaCl; 5mM KCl; 1mM KH_2_PO_4_; 0.2mM MgCl_2_·6H_2_O (Fisher Scientific #M35-500); 0.1mM EGTA; 20mM HEPES; pH 7.4), and incubated on ice for 5min to chelate extracellular Ca^2+^.

Cardiomyocytes were pelleted by centrifugation at 100g for 3 min, the extracellular-like Ca^2+^-free buffer was removed, and the cells were resuspended in permeabilization buffer consisting of intracellular-like medium (120mM KCl; 10mM NaCl; 1mM KH_2_PO_4_; 20mM HEPES; pH 7.2) that had been cleared with Chelex 100 (Bio-Rad # 1422822) to remove trace Ca^2+^, and supplemented with 1X EDTA-free protease inhibitor cocktail (Sigma-Aldrich #4693132001), 120 µg/mL digitonin (Sigma-Aldrich # D141), 3µM thapsigargin (Enzo Life Sciences # BML-PE180-0005), and 5mM succinate (Sigma-Aldrich # S3674). 1 µM Fura-FF (AAT Bioquest #21028) was used to monitor extra-mitochondrial Ca^2+^ and 4.8 µM JC-1 (Enzo Life Sciences #52304) was added at the indicated time to monitor mitochondrial membrane potential (ΔΨ_m_). Permeabilized cardiomyocytes were gently stirred at 37°C in a Delta Ram spectrofluorometer (Photon Technology International) set to record fluorescence at 340nm_ex_, 535nm_em_ and 380nm_ex_, 535nm_em_ for Fura-FF and at 570nm_ex,_ 595nm_em_ for the JC-1 aggregate and 490nm_ex_, 535nm_em_ for the JC-1 monomer. The ratio of Fura-FF fluorescence at 340nm_ex_, 535nm_em_ / 380nm_ex_, 535nm_em_ was plotted to assess extra-mitochondrial Ca^2+^ and the JC-1 570nm_ex_, 595nm_em_ / 490nm_em_, 535nm_ex_ ratio was plotted to assess ΔΨ_m_.

For measurement of _m_Ca^2+^ uptake rate, experiments were performed in the presence of 10µM CGP-37157 (Enzo Life Sciences #BML-CM119-0005) to inhibit _m_Ca^2+^ efflux via NCLX. JC-1 was added to the permeabilized cells after 20sec of baseline recording, and energization of mitochondria was verified by an increase in the ratio of JC-1 aggregate/monomer fluorescence. A bolus of 5µM CaCl_2_ was injected at 350sec, followed by injection of a 10µM bolus of CaCl_2_ at 650sec. The experiment was terminated by addition of 10µM FCCP (Sigma-Aldrich #C2920) to collapse ΔΨ_m_ and release matrix Ca^2+^. 2-3 replicate assay recordings were performed to determine the average Ca^2+^ uptake rate for the 30sec following the peak of the 10µM Ca^2+^ bolus for each mouse.

For measurement of mitochondrial calcium retention capacity, JC-1 was added to permeabilized cells after 20sec of baseline recording. Beginning at 400sec, 10µM boluses of CaCl_2_ were injected every 60sec until spontaneous collapse of ΔΨ_m_ and release of matrix Ca^2+^ to the bath solution, indicative of mitochondrial permeability transition. The experiment was terminated by addition of 10µM FCCP to confirm collapse of ΔΨ_m_. 1-3 replicate assay recordings were performed to determine the average number of Ca^2+^ boluses tolerated prior to ΔΨ_m_ collapse for each mouse.

### Mitochondrial swelling assay

Pelleted mitochondria isolated from the hearts of adult mice as described above were resuspended in fresh IBM2 and centrifuged at 7200g for 12min at 4°C. The supernatant was removed and mitochondrial pellets were resuspended in isolated mitochondria assay buffer (125mM KCl; 20mM HEPES; 2mM MgCl_2_; 2mM KH_2_PO_4_; pH = 7.2). Protein concentration was determined by bicinchoninic acid assay. 300µg of mitochondria/well were added to a 96 well plate in 200µL assay buffer supplemented with 10mM succinate. Absorbance at 540±20nm was measured using a Tecan Infinite M1000 Pro plate reader set at 37°C. A 500µM CaCl_2_ bolus was injected after 2min of baseline recording and mitochondrial swelling was assessed as a decrease in absorbance.

### Extracellular flux assays

Cardiomyocytes were isolated from the hearts of adult mice as described above, and the CaCl_2_ concentration in the stopping buffer was gradually increased to 1mM. Cardiomyocytes were then pelleted by centrifugation at 100g for 3min, and resuspended in DMEM (Corning #90-113-PB) supplemented with 5mM glucose, 4mM L-glutamine (Corning #1-030-RM), 0.1mM sodium pyruvate (Sigma-Aldrich #P8574), 0.2mM BSA-conjugated palmitate (Sigma-Aldrich #A7030; Sigma-Aldrich #P9767), 0.2mM carnitine (Sigma-Aldrich #C0283), pH = 7.4^34^. 1250 live cardiomyocytes/well were plated to a 96-well plate (Agilent #101085-004) coated with 50µg/mL laminin (ThermoFisher #23017-015) and allowed to attach for 1 hour in a CO_2_-free 37°C incubator. A Seahorse XF96 extracellular flux analyzer (Agilent) was used to measure oxygen consumption rate (OCR) at baseline and after sequential additions of 3µM oligomycin (Sigma-Aldrich #O4876); 1.5µM FCCP; and 2µM rotenone (Sigma-Aldrich #R8875) + 2µM antimycin A (Sigma-Aldrich # A8674). Basal, ATP-linked, non-mitochondrial, and maximal respiration; proton leak; and respiratory reserve capacity were calculated as described previously^34^.

### Echocardiography

Left ventricular echocardiography was performed as reported elsewhere^19^ at baseline prior to isoproterenol infusion and at indicated timepoints. In brief, mice were anesthetized with 1.5% isoflurane in 100% oxygen and M-mode images were recorded in the short-axis view suing a Vevo 2100 imaging platform (VisualSonics). Recordings were analyzed with VisualSonics Vevo LAB software (VisualSonics version 3.1.1).

### Isoproterenol infusion

Osmotic minipump implantation surgeries were performed as detailed previously^35^. Mice were anesthetized with 3% isoflurane and an osmotic minipump (Alzet model 2004, #0000298) set to deliver (±)-isoproterenol hydrochloride (Sigma-Aldrich #I5627) dissolved in sterile saline at a dose of 70mg/kg/day for 2 weeks was inserted subcutaneously via a small midline incision in the back. The incision was closed with 5-0 absorbable suture, and mice were administered 40mg/kg of the antibiotic cefazolin (Sandoz #007813450). Mice in the sham group were subjected to the same protocol, but no minipump was inserted.

### Tissue gravimetrics and histology

Hearts were collected 2 weeks after osmotic minipump implantation or sham surgery and massed. Tibia length was measured for normalization of heart mass. The ventricles were rinsed in ice-cold 1X PBS, then divided for further analysis. Ventricle base samples were snap frozen with liquid nitrogen-cooled tongs and stored at −80°C for use in western blotting. Mid-ventricle cross sections were fixed in 10% buffered formalin (EKI #4498), dehydrated, embedded in paraffin, and cut to 7µm sections and mounted on glass slides. Heart sections were labelled with TRITC-conjugated wheat germ agglutinin (Sigma-Aldrich #L5266) at 100µg/mL to outline each cell, and coverslips were mounted using ProLong Gold Antifade Mountant with DAPI (Invitrogen #P36935). Slides were imaged and cardiomyocyte cross-sectional area measured as described previously^35^. Lung wet mass was measured at the time of collection, and lung dry mass was measured after drying the tissue at 37°C for 48 hours for assessment of lung edema.

### *In vitro* assessment of reactive oxygen species generation

Isolated adult mouse cardiomyocytes were prepared and plated as described for extracellular flux assays above at 5000 live cells/well to clear-bottomed, black-walled 96-well plates (Greiner Bio-One #655090) coated with 50µg/mL laminin. Cells were allowed to attach for 1 hour. The plates were then changed to fresh media supplemented with 10µM ionomycin (Cayman Chemical #11932) or 2µM antimycin A or vehicle control and incubated at 37°C for 1hr. Dihydroethidium (DHE) (ThermoFisher #D11347) was added to a final concentration of 5µM for the final 30 minutes of this incubation period. After 1hr total treatment time, the fluorescence of oxidized DHE was measured at 500nm_ex_, 580nm_em_ using a Tecan Infinite M1000 Pro plate reader.

### *In vitro* cell death assays

Isolated adult mouse cardiomyocytes were plated at 1250 live cells/well to laminin-coated, clear-bottomed, black-walled 96-well plates as for the assessment of reactive oxygen species generation described above. After 1hr, the plates were changed to fresh media supplemented with 30µM propidium iodide (ThermoFisher #P1304MP) and with 10-50µM ionomycin or 2mg/L (±)-isoproterenol hydrochloride or vehicle control and incubated at 37°C for 1hr. Propidium iodide fluorescence was then measured at 530nm_ex_, 645nm_em_ on a Tecan Infinite M1000 Pro plate reader.

### Evans blue dye labeling

After 2 weeks of isoproterenol infusion, mice were injected I.P. with a sterile 1% (w/v) solution of Evans blue dye (EBD) (Sigma-Aldrich #E2129) dissolved in 1X PBS, at a volume of 1% of body mass^36^. Hearts were collected 18hrs later as described above, and mid-ventricle cross sections were placed in Tissue-Tek O.C.T. Compound (Sakura #4583) and frozen in liquid nitrogen-cooled isopentane. Samples were cut to 5µm cross sections, mounted on glass slides, and labelled with wheat germ agglutinin-AlexaFluor 488 conjugate (ThermoFisher #W11261) at 100µg/mL to outline each cell. Coverslips were mounted using ProLong Gold Antifade Mountant with DAPI. Slides were imaged on a Nikon Eclipse Ti-E fluorescence microscope. The average percentage of cardiomyocytes stained with EBD across 6 10x fields of view per heart was quantified.

### Statistics

All data are presented as mean ± S.E.M. unless otherwise indicated. Statistical analyses were carried out with Prism 6.0 (GraphPad Software). A two-tailed t-test was used for direct comparisons between two groups. Welch’s correction was used in cases of unequal variance. Kaplan Meier survival curves were compared using the log-rank (Mantel-Cox) test. Longitudinal echocardiographic studies were analyzed using 2-way ANOVA. Dunnett’s post-hoc analysis was used for comparison to a single control, and Sidak’s post-hoc analysis was used for comparisons across multiple groups. Grouped endpoint data were analyzed by 2-way ANOVA with Sidak’s post-hoc analysis. Dose-response curves were evaluated by non-linear regression using a least squares ordinary fit, and corresponding EC_50_ values were compared by extra-sum-of squares F-test. For all analyses, *P* values less than 0.05 were considered significant.

## RESULTS

### Development and validation of a genetic mouse model of conditional cardiomyocyte-specific MCU overexpression

To investigate the contribution of mtCU-dependent _m_Ca^2+^ uptake to cardiac stress responses *in vivo*, we developed a gain-of-function flox-stop mouse model (“flox-stop-MCU”) allowing for Cre-dependent, temporally controlled expression of a human *MCU* transgene in adult mouse cardiomyocytes when crossed to mice with the α-myosin heavy chain-MerCreMer (αMHC-MCM; “MCM”) allele (**Fig. 1A**). We selected a founder line with no MCU transgene expression in the absence of Cre, and with robust MCU transgene expression when crossed to mice with the constitutive, cardiomyocyte-restricted αMHC-Cre allele (**Supplemental Fig. S1A**). Treatment of αMHC-MCM x flox-stop-MCU (“MCU-Tg”) mice with tamoxifen increased total cardiomyocyte MCU protein content within 1-wk after the first tamoxifen dose (**Fig. 1B**). Fast protein size-exclusion liquid chromatography (FPLC) of lysates of cardiac mitochondria isolated from adult mice verified an approximate 20-fold increase in total MCU content in MCU-Tg mitochondria (**Fig. 1 C-E**). Overexpressed MCU in MCU-Tg cardiac mitochondria distributed into protein fractions ranging up to ∼800kD, similar to the molecular weight distribution of endogenous mouse MCU (**Fig. 1 C-D**). These data suggest that overexpressed MCU assembles with the correct stoichiometry with other mtCU complex components to form intact, high-molecular weight uniporter channels within cardiac mitochondria. Despite robust mitochondrial MCU overexpression in MCU-Tg hearts, we did not detect any significant change in mRNA transcript expression of other mtCU components, nor of genes that mediate _m_Ca^2+^ efflux (**Supplemental Fig. S1 B-I**). This result also suggests that post-transcriptional regulatory mechanisms are likely influencing uniporter assembly and maintenance.

**Figure 1:**
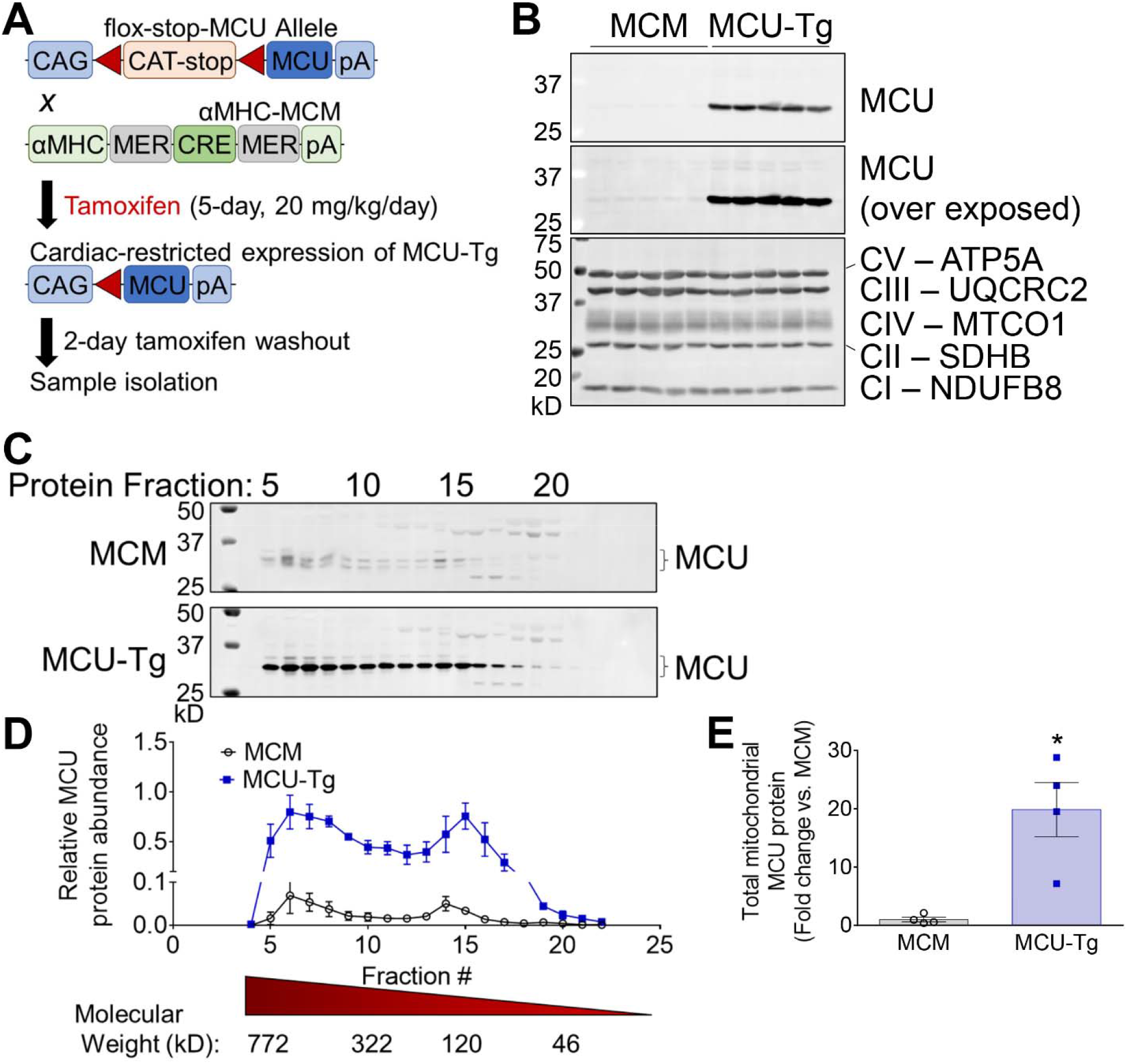
Mouse model of inducible, adult cardiomyocyte-specific MCU transgene expression. **A**) Genetic approach for conditional overexpression of MCU transgene in adult mouse cardiomyocytes. **B**) Western blots for tamoxifen-inducible overexpression of MCU in adult cardiomyocytes isolated from αMHC-MCM (MCM) and αMHC-MCM x flox-stop-MCU (MCU-Tg) mice. Total OXPHOS complexes I-V are shown as a mitochondrial loading control. Corresponding full-length blots are shown in Supplemental Fig. S2. (*n*=5 mice/genotype). **C**) Western blots for MCU of fast protein size-exclusion liquid chromatography fractions of cardiac mitochondria isolated from MCM and MCU-Tg hearts. Brackets indicate quantified mature, mitochondrial targeting sequence-cleaved MCU. Corresponding full-length blots are shown in Supplemental Fig. S2. **D**) Relative distribution of MCU in size exclusion chromatography fractions of isolated cardiac mitochondria. (*n*=4 mice per genotype). **E**) Quantification of total MCU protein across FPLC fractions from MCM and MCU-Tg cardiac mitochondria. Data analyzed by unpaired, two-tailed *t-*test. \**p*<0.05. (*n*=4 mice/genotype).

We next assessed the functional impact of increased cardiomyocyte MCU content by examining the effect of MCU overexpression on _m_Ca^2+^ handling. Evaluation of acute _m_Ca^2+^ uptake in isolated, permeabilized adult mouse cardiomyocytes 1-wk after the start of tamoxifen treatment revealed an accelerated rate of _m_Ca^2+^ uptake in MCU-Tg cells (**Fig. 2 A-B**). This indicates that overexpression of MCU alone is sufficient to increase uniporter function in adult mouse cardiomyocytes. Consistent with enhanced net _m_Ca^2+^ uptake, MCU-Tg cardiomyocytes required the addition of fewer successive Ca^2+^ boluses to reach mitochondrial permeability transition, as indicated by a collapse of mitochondrial membrane potential, ΔΨ_m_, and release of Ca^2+^ from the mitochondrial matrix (**Fig. 2 C-D**). Isolated MCU-Tg cardiac mitochondria also exhibited increased swelling in response to a single 500-µM Ca^2+^ bolus (**Fig. 2 E-G**), reflecting an increased propensity for _m_Ca^2+^ uptake, which triggers subsequent permeability transition.

**Figure 2:**
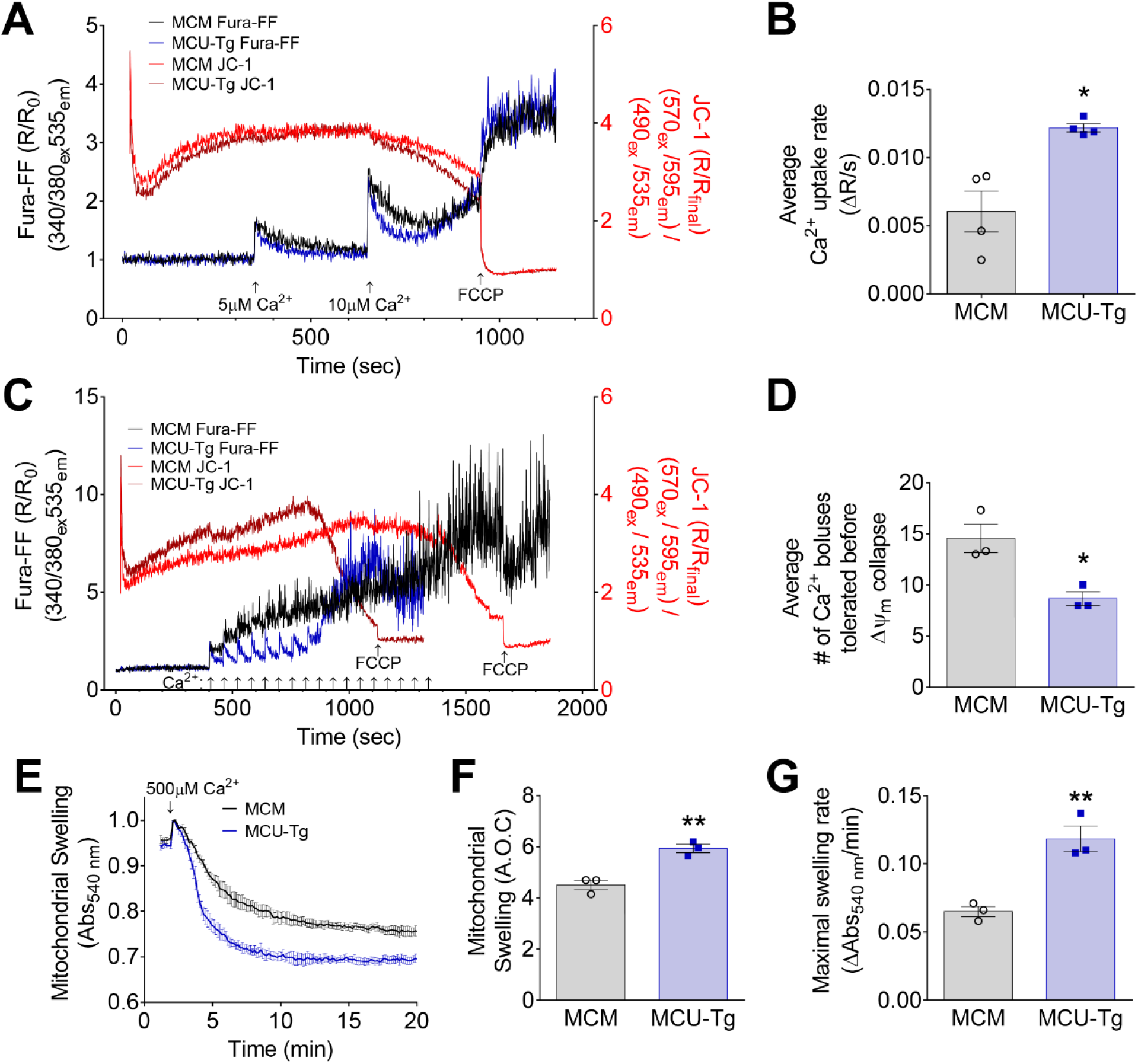
Transgenic MCU overexpression increases _m_Ca^2+^ uptake in adult mouse cardiomyocytes. **A**) Mean traces showing _m_Ca^2+^ uptake in permeabilized adult αMHC-MCM (MCM) and αMHC-MCM x flox-stop-MCU (MCU-Tg) mouse cardiomyocytes in response to 5 or 10µM Ca^2+^ bolus delivered at indicated timepoints. Cardiomyocytes were isolated 1-wk after the start of tamoxifen treatments and permeabilized with digitonin. Measurements were performed in the presence of CGP-37157 to inhibit _m_Ca^2+^ efflux through NCLX and thapsigargin to inhibit Ca^2+^ uptake through SERCA. Fura-FF fluorescence represents extra-mitochondrial, bath Ca^2+^ content. JC-1 fluorescence represents mitochondrial membrane potential, ΔΨ_m_. Traces are mean of 11 recordings/genotype. **B**) Quantification of average _m_Ca^2+^ uptake rate for each mouse, measured over the first 30s following the peak of the 10µM Ca^2+^ bolus. Data analyzed by unpaired, two-tailed *t*-test. \**p*<0.05. (*n*=4 mice per genotype). **C**) Ca^2+^ retention capacity (CRC) assay in permeabilized adult mouse cardiomyocytes. 10μM Ca^2+^ boluses were added every 60s as indicated. Measurements were made in the presence of thapsigargin. Traces are mean of 6 recordings from MCM mice and 7 recordings from MCU-Tg mice. **D**) Average number of 10µM Ca^2+^ boluses tolerated in CRC assay before Dy_m_ (mitochondrial membrane potential) collapse for each mouse. Data analyzed by unpaired, two-tailed *t*-test. \**p*<0.05. (*n*=3 mice/genotype). **E**) Assay showing swelling (decrease in absorbance) of isolated cardiac mitochondria in response to addition of a 500µM Ca^2+^ bolus. Traces are mean ± S.E.M. (*n*=3 mice/genotype). Quantification of mitochondrial swelling (area over the curve, A.O.C.) (**F**) and maximal swelling rate (**G**) in isolated cardiac mitochondria in response to addition of a 500µM Ca^2+^ bolus. Data analyzed by unpaired, two-tailed *t*-test. \*\**p*<0.01. (*n*=3 mice/genotype).

We then examined mitochondrial metabolism to assess whether the increased capacity for _m_Ca^2+^ uptake observed with MCU overexpression had physiological consequences in intact adult cardiomyocytes. Inhibitory phosphorylation of the pyruvate dehydrogenase (PDH) E_1_α subunit at serine 293 was decreased in MCU-Tg cardiomyocytes (**Fig. 3 A-B**), consistent with increased _m_Ca^2+^ uptake leading to net _m_Ca^2+^ accumulation and increased activity of the _m_Ca^2+^-sensitive PDH phosphatase. MCU overexpression increased basal and ATP-linked oxygen consumption in isolated intact adult cardiomyocytes (**Fig. 3 C-D**), consistent with increased activity of PDH and TCA cycle dehydrogenases. These findings show that mitochondrial metabolism is increased in intact MCU-Tg cardiomyocytes, in support of the notion that basal _m_Ca^2+^ signaling is elevated in these cells under homeostatic conditions. Together, our *in vitro* results validate the MCU-Tg mouse model as a genetic tool for enhanced _m_Ca^2+^ uptake, even in normal physiological contexts.

**Figure 3:**
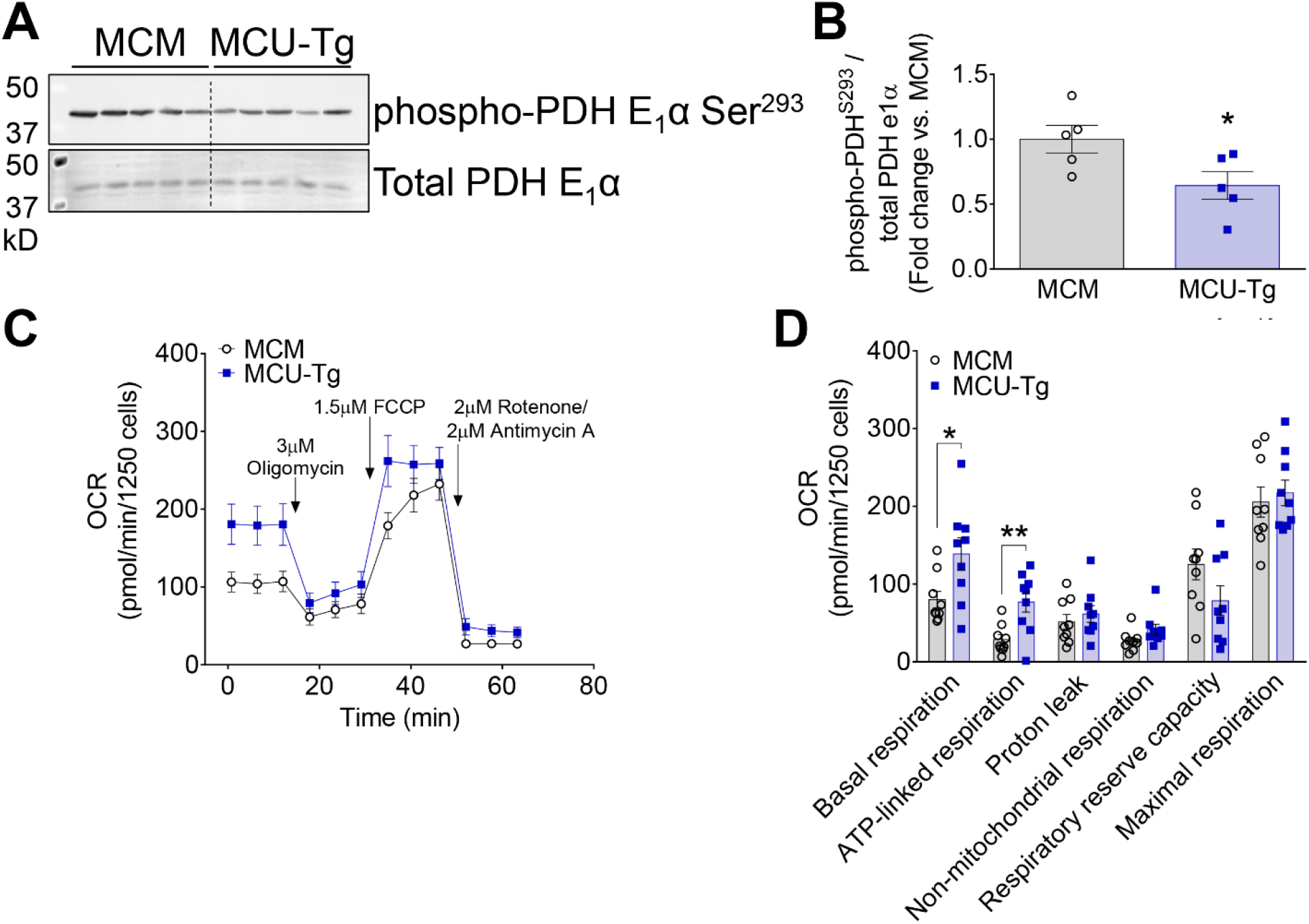
MCU overexpression increases basal mitochondrial respiration in adult mouse cardiomyocytes. **A**) Western blots for pyruvate dehydrogenase (PDH) phosphorylation in adult cardiomyocytes isolated from αMHC-MCM (MCM) and αMHC-MCM x flox-stop-MCU (MCU-Tg) mice 1-wk after the administration of tamoxifen. Blots were performed using the same cardiomyocyte samples as shown in Fig. 1B. Corresponding full-length blots are shown in Supplemental Fig. S3. **B**) Semi-quantification of inhibitory PDH E1α S293 phosphorylation. Data analyzed by unpaired, two-tailed *t*-test. \**p*<0.05. (*n*=5 mice per genotype). **C**) Extracellular flux analysis of oxygen consumption rate (OCR) in isolated adult mouse cardiomyocytes. Traces represent mean ± S.E.M.; 9 mice/genotype. **D**) Quantification of basal respiration, ATP-linked respiration, proton leak, non-mitochondrial respiration, respiratory reserve capacity, and maximal respiration. Data analyzed by unpaired, two-tailed *t*-test. \**p*<0.05, \*\**p*<0.01. (*n*=9 mice/genotype).

### mtCU-dependent _m_Ca^2+^ uptake is required for contractile responsiveness to β-adrenergic stimulation *in vivo*, but drives cardiac maladaptation during prolonged stress

Following validation of the novel gain-of-function model, we compared mice with adult cardiomyocyte-specific MCU overexpression and mice with inducible, adult cardiomyocyte-specific *Mcu* deletion (αMHC-MCM x Mcu*^fl/fl^*; *Mcu*-cKO)^19^ to address the question of how mtCU-dependent _m_Ca^2+^ uptake impacts the heart’s immediate and long-term adaptation to a sustained increase in workload. After 5 days of tamoxifen administration, mice were allowed 16 days of tamoxifen washout to allow time for turnover of MCU protein (**Fig. 4A** and **Supplemental Fig. S4**). Three wks after the first tamoxifen dose, MCM, *Mcu*-cKO, and MCU-Tg mice were implanted with osmotic minipumps to deliver isoproterenol (70mg/kg/day) (**Fig. 4A**). We noted a slight decrease in survival among *Mcu*-cKO and MCU-Tg mice throughout 14 days of isoproterenol infusion, although this did not reach statistical significance (**Fig. 4B**). Isoproterenol infusion significantly increased cardiac contractility within 2 days of the start of isoproterenol infusion in MCM controls and this effect was abrogated in *Mcu*-cKO mice (**Fig. 4 C-E**). Despite this lack of contractile responsiveness to isoproterenol, we did not observe any detrimental effect of adult cardiomyocyte-specific *Mcu* deletion on contractile function under baseline conditions (day 0), nor any measurable decline in cardiac contractility of *Mcu*-cKO hearts over 14-day isoproterenol infusion (**Fig. 4 C-E**). Baseline cardiac function and the initial increase in cardiac contractility at 2 days of isoproterenol infusion were largely preserved in MCU-Tg hearts (**Fig. 4 C-E**). In contrast, by 7-14 days of isoproterenol infusion, MCU overexpression resulted in a significant decline in contractile function compared both to baseline and to MCM controls (**Fig. 4 C-E**). These findings support the notion that increased capacity for _m_Ca^2+^ uptake through the mtCU is not deleterious to cardiomyocyte function unless combined with an additional stressor, such as elevated cytosolic Ca^2+^ cycling due to chronic β-adrenergic stimulation. However, with sufficient duration of persistent stress signaling and intracellular Ca^2+^ (_i_Ca^2+^) load, the resulting ongoing increase in _m_Ca^2+^ uptake becomes detrimental to the overall contractile function of the heart.

**Figure 4:**
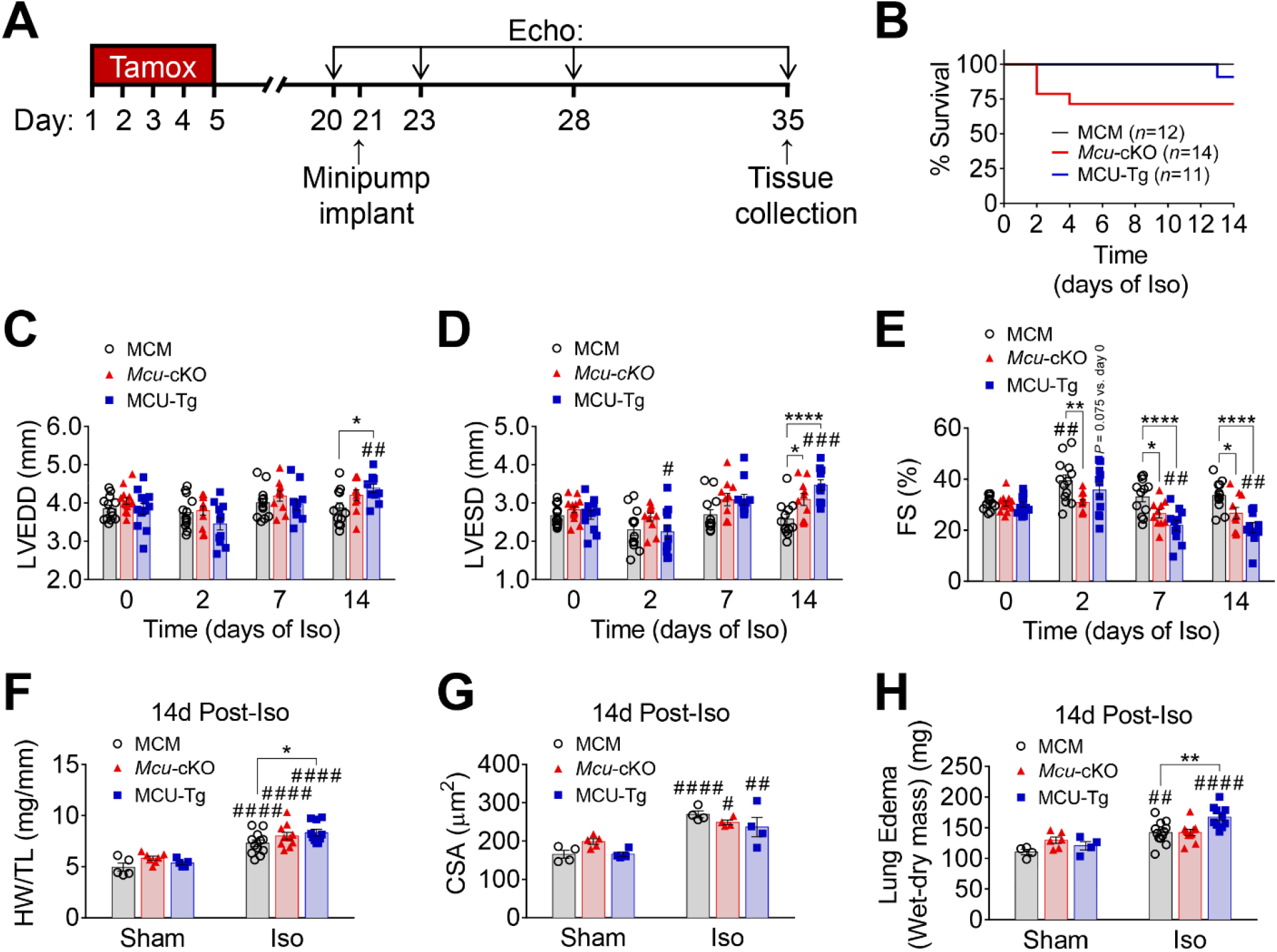
Cardiomyocyte *Mcu* deletion prevents isoproterenol-induced increase in cardiac contractility, while MCU overexpression predisposes to isoproterenol-induced contractile dysfunction. **A)** Experimental timeline of tamoxifen administration, isoproterenol minipump implant surgery, and *in vivo* functional analysis in αMHC-MCM (MCM), αMHC-MCM x Mcu*^fl/fl^ (Mcu*-cKO), and αMHC-MCM x flox-stop-MCU (MCU-Tg) mice. **B)** Kaplan-Meier survival curves of MCM, Mcu-cKO, and MCU-Tg mice. Mice/group at the start of the study is indicated in parentheses. Data analyzed by log-rank (Mantel-Cox) test. Left ventricular end-diastolic dimension (LVEDD) **(C)**, end-systolic dimension (LVESD) **(D)**, and percent fractional shortening (%FS) **(E)** over 14-days of Iso infusion. Data analyzed by 2-way ANOVA with Dunnett’s post-hoc test. \**p*<0.05, \*\**p*<0.01, \*\*\*\**p*<0.0001 vs. MCM; ^#^*p*<0.05, ^##^*p*<0.01, ^###^*p*<0.001 vs. day 0. (*n*=12-13 MCM, 10-13 *Mcu*-cKO, 10-12 MCU-Tg mice). Heart weight-to-tibia length (HW/TL) ratio **(F)** (Sham: *n*=5 MCM, 7 *Mcu*-cKO, 5 MCU-Tg mice; Iso: *n*= 12 MCM, 10 *Mcu*-cKO, 10 MCU-Tg mice); cardiomyocyte cross sectional area (CSA) **(G)** (Sham: *n*=4 MCM, 5 *Mcu*-cKO, 4 MCU-Tg mice; Iso: *n*=4 mice/genotype); and lung edema **(H)** (Sham: *n*=4 MCM, 6 *Mcu*-cKO, 4 MCU-Tg mice; Iso: *n*=12 MCM, 10 *Mcu*-cKO, 10 MCU-Tg mice) at 14-day endpoint. Data analyzed by 2-way ANOVA with Sidak’s post-hoc test. \**p*<0.05, \*\**p*<0.01 vs. MCM; ^#^*p*<0.05, ^##^*p*<0.01, ^####^*p*<0.0001 vs. Sham.

Despite its disparate effects on the contractile performance of control MCM versus *Mcu*-cKO hearts, 14-days of isoproterenol infusion increased the heart weight-to-tibia length (HW/TL) ratio and cardiomyocyte cross-sectional area (CSA) to a similar extent in both genotypes (**Fig. 4 F-G**). The increase in HW/TL after 14-days of isoproterenol was exaggerated in MCU-Tg hearts (**Fig. 4F**), although the increase in cardiomyocyte cross-sectional area was similar between all genotypes (**Fig. 4G**). Lung edema was also exaggerated in MCU-Tg mice (**Fig. 4H**), in agreement with their worsened contractile function compared to controls.

### Increased mtCU activity enhances ROS production and sensitizes cardiomyocytes to cell death *in vitro*

To explore the mechanisms by which increased MCU expression contributes to contractile dysfunction in response to sustained sympathetic stress, we tested the responses of isolated adult cardiomyocytes to cellular Ca^2+^ stress *in vitro*. MCU-Tg cardiomyocytes demonstrated a tendency for increased cellular reactive oxygen species (ROS) production, indicated by a greater increase in DHE fluorescence, upon incubation with the Ca^2+^ ionophore ionomycin (**Fig. 5A**). MCU-Tg cardiomyocytes also exhibited exaggerated ROS production in response to incubation with the respiratory complex III inhibitor, antimycin A (**Fig. 5B**). These results suggest that increased _m_Ca^2+^ uptake contributes to potentially deleterious ROS production. Consistent with this model, MCU-Tg cardiomyocytes displayed increased sensitivity to cell death induced by the Ca^2+^ ionophore, ionomycin (**Fig. 5C**), and a trend towards increased membrane rupture during acute 1-hr isoproterenol stimulation (**Fig. 5D**). Together, these findings indicate that enhanced _m_Ca^2+^ uptake in MCU-Tg cardiomyocytes renders them more susceptible to oxidative stress and cell death when subjected to stimuli that increase cytosolic Ca^2+^ concentration.

**Figure 5:**
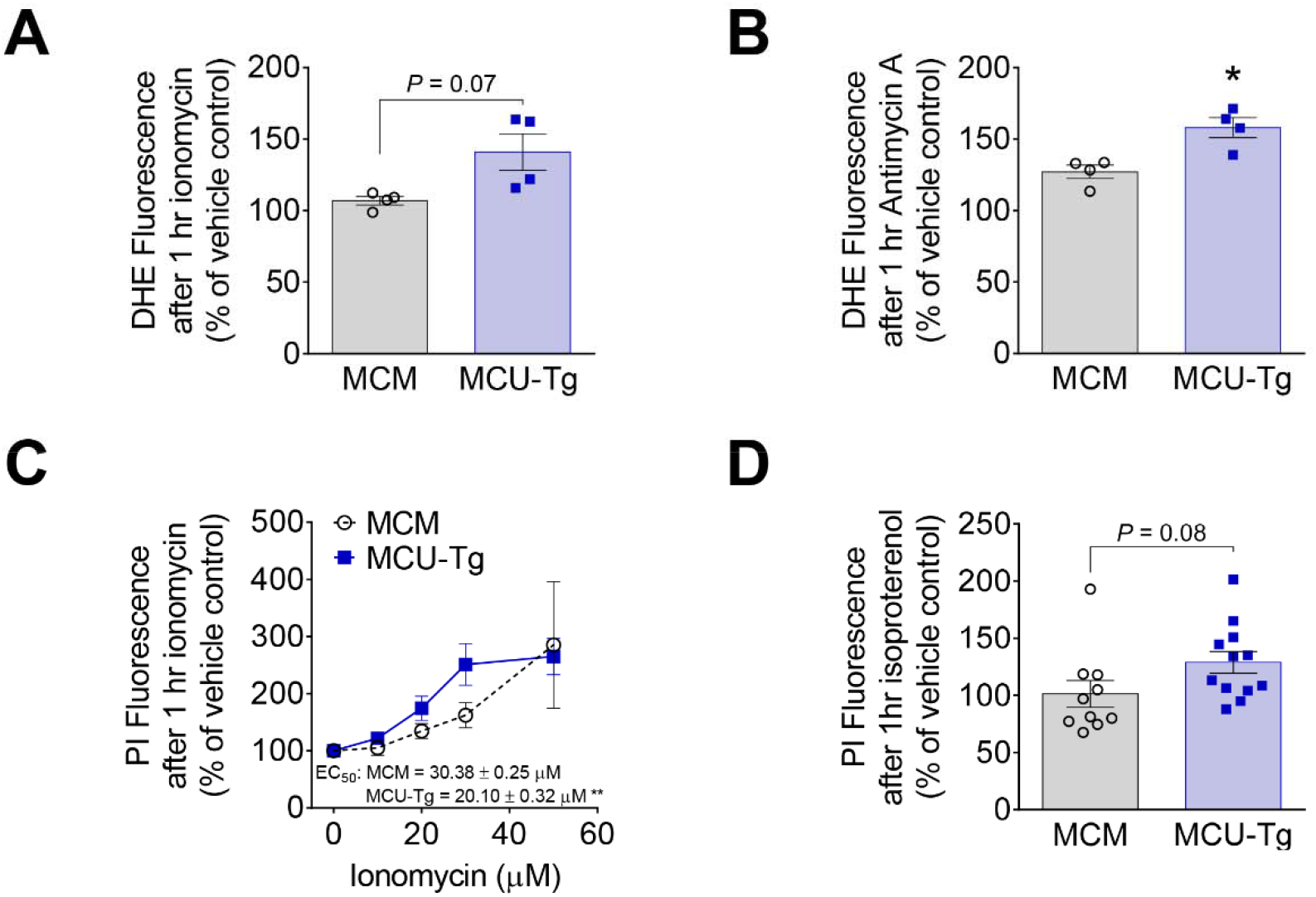
Increased _m_Ca^2+^ uptake increases Ca^2+^-induced reactive oxygen species (ROS) production and cell death in adult cardiomyocytes. **A)** ROS generation as indicated by dihydroethidium (DHE) fluorescence in isolated adult αMHC-MCM (MCM) and αMHC-MCM x flox-stop-MCU (MCU-Tg) mouse cardiomyocytes incubated with the Ca^2+^ ionophore, ionomycin. Data analyzed by unpaired, two-tailed *t*-test. (*n*=4 mice/genotype). **B)** ROS generation in isolated adult mouse cardiomyocytes incubated with the respiratory complex III inhibitor, antimycin A. Data analyzed by unpaired, two-tailed *t*-test. \**p*<0.05. (*n*=4 mice/genotype). **C)** Cell death as indicated by increased propidium iodide (PI) fluorescence in isolated adult mouse cardiomyocytes incubated with increasing doses of ionomycin. Best-fit EC_50_ values compared by extra-sum-of squares F-test. \*\**p*<0.01 vs. MCM. (*n*=10 MCM mice, 12 MCU-Tg mice). **D)** Cell death as indicated by an increase in PI fluorescence in isolated adult mouse cardiomyocytes incubated with isoproterenol. Data analyzed by unpaired, two-tailed *t*-test. (*n*=10 MCM mice, 12 MCU-Tg mice).

### Isoproterenol-induced contractile dysfunction and cardiomyocyte death in MCU-Tg mice is not rescued by genetic inhibition of the mPTP

Mitochondrial Ca^2+^ overload is a classical stimulus for the generation of ROS and mPTP opening leading to necrotic cell death^2, 37, 38^, and oxidative stress itself can trigger mitochondrial permeability transition (mPT)^31, 39, 40^. Cardiomyocyte dropout due to apoptosis and/or necrosis has been proposed as a mechanism underlying the decline in contractile function in heart failure^41, 42^. Further, disruption of the mPTP is sufficient to rescue the lethal phenotype induced by rapid _m_Ca^2+^-overload resulting from inducible NCLX deletion in adult cardiomyocytes^25^. We therefore hypothesized that the progressive contractile dysfunction we observed in MCU-Tg mice with isoproterenol infusion was caused by cardiomyocyte dropout due to mitochondrial Ca^2+^ overload, activation of the mPTP, and subsequent necrosis. We crossed MCU-Tg and MCM mice to *Ppif* ^-/-^ mice lacking the mPTP regulator cyclophilin D (CypD) to test whether genetic inhibition of the mPTP is sufficient to rescue isoproterenol-induced contractile dysfunction of MCU-Tg hearts. Western blotting confirmed loss of CypD protein in *Ppif*^-/-^ mice, and persistence of MCU overexpression in MCU-Tg hearts on both wild-type and *Ppif^-^*^/-^ backgrounds (**Fig. 6A; Supplemental Fig. S6A**). Interestingly, MCU overexpression also drove an increase in cardiac protein expression of the core mtCU component EMRE in mice on both wild-type and *Ppif*^-/-^ backgrounds (**Fig. 6A; Supplemental Fig. S6B**), even though it did not affect levels of EMRE transcript **(Supplemental Fig. S1B**). Increased EMRE expression in MCU-Tg hearts persisted even after 14-days of isoproterenol infusion, but the extent of this increase in EMRE was somewhat attenuated with isoproterenol infusion in MCU-Tg x *Ppif*^-/-^ hearts. (**Fig. 6A; Supplemental Fig. S6B**). Cardiac expression of the mtCU gatekeeper MICU1 did not differ among genotypes under control conditions, but upon chronic isoproterenol infusion, MICU1 protein expression was decreased in MCU-Tg hearts and tended to be downregulated to a similar extent in MCU-Tg x *Ppif*^-/-^ hearts (**Fig. 6A; Supplemental Fig. S6C**). These results suggest that modest post-translational compensatory changes to regulators of uniporter function occur in the context of MCU overexpression, and that these adaptations are also responsive to increased _m_Ca^2+^ loading during isoproterenol stimulation.

**Figure 6:**
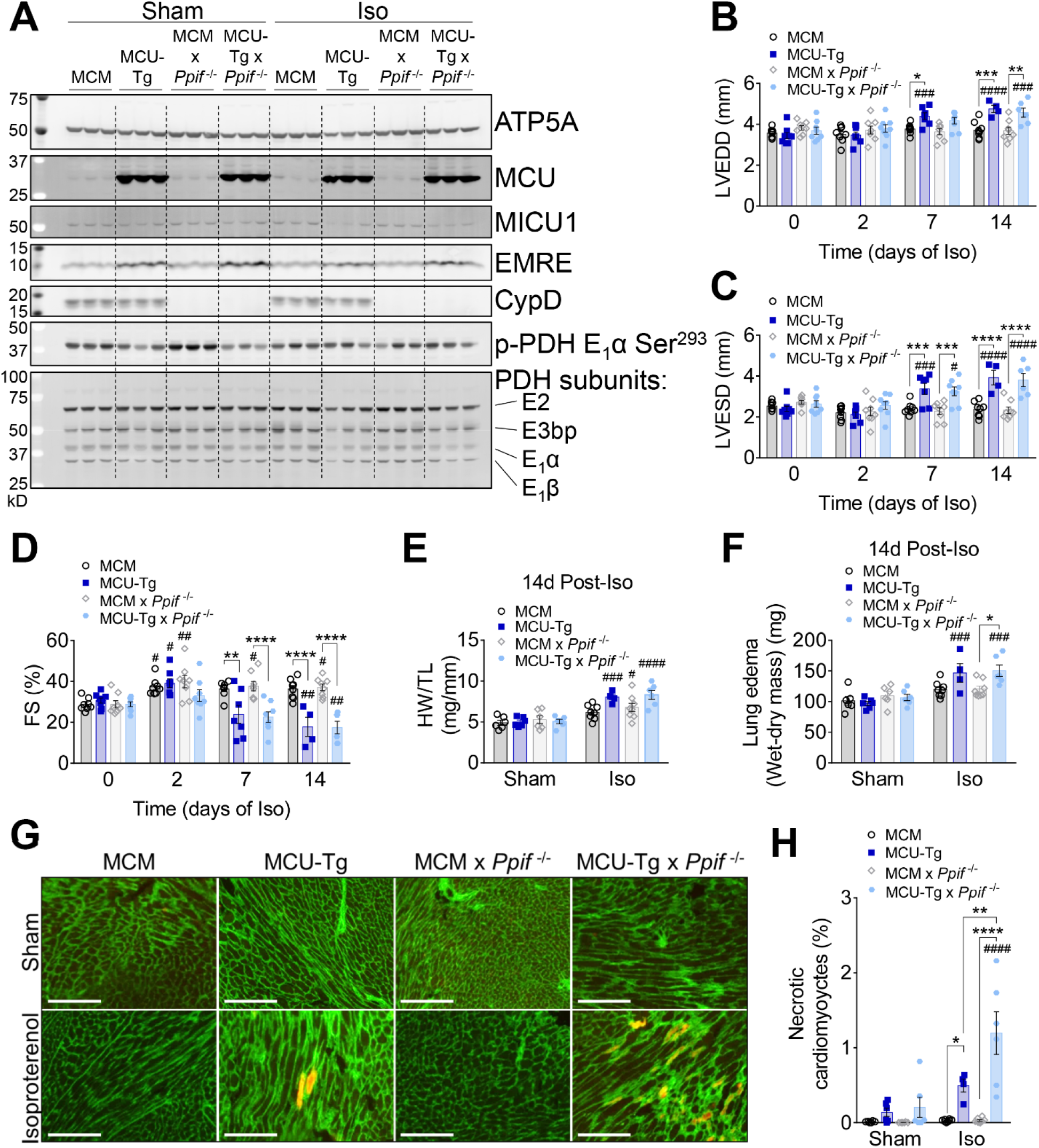
Genetic inhibition of the mPTP (*Ppif*^-/-^) does not prevent contractile dysfunction, remodeling, or cardiomyocyte death in MCU-Tg hearts during chronic isoproterenol infusion. **A)** Western blots confirming tamoxifen-inducible overexpression of MCU in MCU-Tg hearts and constitutive deletion of cyclophilin D (CypD) in *Ppif*^-/-^ hearts. Expression of mtCU components EMRE and MICU1, and pyruvate dehydrogenase (PDH) phosphorylation and subunit expression were also examined. Hearts were collected after 14-days of isoproterenol (Iso) infusion. ATP5A is shown as a mitochondrial loading control. Corresponding full-length blots are shown in Supplemental Fig. S5. Left ventricular end-diastolic dimension (LVEDD) **(B)**, end-systolic dimension (LVESD) **(C)**, and percent fractional shortening (%FS) **(D)** over 14 days of Iso infusion. Data analyzed by 2-way ANOVA with Sidak’s post-hoc test. \**p*<0.05, \*\**p*<0.01, \*\*\**p*<0.001, \*\*\*\**p*<0.0001 between genotypes; ^#^*p*<0.05, ^##^*p*<0.01, ^###^*p*<0.001, ^####^*p*<0.0001 vs. day 0. (*n*=8 MCM; 4-8 MCU-Tg; 8 MCM x *Ppif*^-/-^; and 6-8 MCU-Tg x *Ppif*^-/-^ mice). Heart weight-to-tibia length (HW/TL) ratio **(E)** and lung edema **(F)** at 14-day endpoint. Data analyzed by 2-way ANOVA with Sidak’s post-hoc test. \**p*<0.05 between genotypes; ^#^*p*<0.05, ^###^*p*<0.001, ^####^*p*<0.0001 vs. Sham. (Sham: *n*=6 mice/genotype; Iso: *n*=8 MCM, 4 MCU-Tg, 8 MCM x *Ppif*^-/-^, and 6 MCU-Tg x *Ppif*^-/-^ mice). **G)** Wheat germ agglutinin (green) and Evans blue dye (EBD) (red) staining in the myocardium at 14-day endpoint. Scale bars = 200 μm. **H)** Percentage of cardiomyocytes stained with EBD as an index of membrane compromise and necrosis. Data analyzed by 2-way ANOVA with Sidak’s post-hoc test. \**p*<0.05, \*\**p*<0.01, \*\*\*\**p*<0.0001 between genotypes; ^####^*p*<0.0001 vs. Sham. (Sham: *n*=6 mice/genotype; Iso: *n*=8 MCM, 4 MCU-Tg, 8 MCM x *Ppif*^-/-^, and 6 MCU-Tg x *Ppif*^-/-^ mice).

Cyclophilin D deletion failed to rescue the contractile dysfunction observed in MCU-Tg hearts after 7-14 days of isoproterenol infusion (**Fig. 6 B-D**). CypD deletion also failed to attenuate the enhanced cardiac hypertrophy and lung edema observed in MCU-Tg mice with chronic isoproterenol infusion (**Fig. 6 E-F**). Therefore, we injected mice with Evans blue dye prior to heart collection to label necrotic cardiomyocytes and examine whether mPTP inhibition effectively limited cardiomyocyte necrosis. MCU-Tg hearts displayed significantly increased cardiomyocyte necrosis only after sustained isoproterenol stimulation (**Fig. 6 G-H**), consistent with their contractile phenotype. CypD deletion failed to decrease, and rather increased the extent of isoproterenol-induced cardiomyocyte necrosis in MCU-Tg hearts (**Fig. 6 G-H**). This finding aligns with the failure of CypD deletion to protect MCU-Tg hearts from isoproterenol-induced contractile dysfunction. We incidentally noted that total protein expression of the pyruvate dehydrogenase (PDH) E_1_α subunit was reduced with isoproterenol infusion in MCU-Tg hearts and tended to be downregulated with isoproterenol infusion in MCU-Tg x *Ppif*^-/-^ hearts (**Fig. 6A and Supplemental Fig. S6D**). Interestingly, the downregulation of PDH E_1_α subunit expression correlated strongly with the diminished left ventricular %FS observed in MCU-Tg and MCU-Tg x *Ppif* ^-/-^ hearts after 14-days of isoproterenol infusion (**Supplemental Fig. S6E**).

## DISCUSSION

Since the genetic identification of the mitochondrial calcium uniporter protein, MCU^3, 4^, research by numerous laboratories has provided insight into the molecular mechanisms by which the mitochondrial calcium uniporter channel assembles and is regulated in order to control _m_Ca^2+^ uptake in the face of fluctuating cytosolic Ca^2+^ levels. Despite growing consensus around the physiological roles of acute _m_Ca^2+^ uptake through the mtCU for physiological cardiac responses, and its contribution to acute ischemic cardiac injury, whether uniporter-dependent _m_Ca^2+^ uptake plays any causative role in the heart’s functional responses to a sustained increase in cardiac workload has remained elusive. This question is particularly relevant for our understanding of the development and progression of forms of heart failure caused by chronic stresses, such as pressure-and/or neurohormonal overload in patients with hypertension. Our results in mice with adult, cardiomyocyte-specific loss-or gain-of MCU function reveal a strict requirement for uniporter-dependent _m_Ca^2+^ uptake in enhancing cardiac contractility throughout the first few days of chronic catecholaminergic stimulation and support a causative role for persistent cardiomyocyte _m_Ca^2+^ loading in driving eventual functional decompensation in response to sustained stress signaling. Interestingly, these findings differ from recent investigations of the role of uniporter-dependent _m_Ca^2+^ uptake in the heart’s response to chronic stress^20, 43^. These discrepancies warrant a more nuanced consideration of the effects of _m_Ca^2+^ loading in general, and of _m_Ca^2+^ uptake specifically through the mtCU versus through alternative pathways, in the development of heart disease. They also raise concern around the potential efficacy and appropriate point of therapeutic intervention for strategies that aim to increase _m_Ca^2+^ uptake to mitigate the progression of heart failure by enhancing mitochondrial energetics. Finally, our findings support a role for persistent _m_Ca^2+^ loading in initiating cardiomyocyte death even independent of classical cyclophilin D-dependent activation of mitochondrial permeability transition, and highlight the need for further investigation into alternative mechanisms that mediate the pathological consequences of _m_Ca^2+^ overload in the diseased heart.

We describe a novel genetic mouse model of tamoxifen-inducible overexpression of an untagged, human MCU transgene that will aid future investigations of tissue-specific mtCU function in the physiology and disease. Overexpressed MCU incorporated into high-molecular weight protein fractions of similar molecular weight to those containing endogenous mouse MCU, suggesting that overall native mtCU subunit stoichiometry and channel composition is maintained when the MCU content of cardiomyocyte mitochondria is increased (**Fig. 1**). Indeed, we found that transgenic MCU overexpression was associated with increased EMRE protein expression (**Fig. 6A**; **Supplemental Fig. S6 A-B**). The concerted upregulation of EMRE in MCU-Tg hearts occurs at the post-transcriptional level, as MCU overexpression did not alter levels of cardiac *Emre* transcript levels (**Supplemental Fig. S1B**). The fact that MICU1 protein expression was not also increased alongside increased MCU and EMRE content in MCU-Tg hearts (**Fig. 6A; Supplemental Fig. S6C**) raises the question of whether the MICU1/MCU ratio may be slightly diminished in this model as compensation for overall increases in uniporter density. This finding may also support a more minor role for MICU1 in the regulation of cardiomyocyte mtCU function, consistent with the conclusion by the Lederer and Boyman group that there is minimal gating of the mtCU in the mature heart^44^. Increased _m_Ca^2+^ uptake in MCU-Tg hearts was well tolerated under unstressed conditions, at least through the 5-wks following induction of MCU overexpression, as we did not observe any signs of pathological cardiac remodeling or evidence for cardiac dysfunction in the absence of isoproterenol stimulation (**Figs. 4** and **6**).

14-day infusion with high-dose isoproterenol tended to increase mortality in *Mcu*-cKO mice with adult cardiomyocyte-specific disruption of uniporter function (**Fig. 4B**). All deaths of *Mcu*-cKO mice occurred within the first few days of isoproterenol infusion, suggesting a critical protective role for mtCU-dependent _m_Ca^2+^ uptake at the onset of a sudden increase in cardiac workload, cellular Ca^2+^ cycling, and energetic demand. This pattern recapitulates our earlier finding of increased mortality at the onset of high-dose angiotensin II + phenylephrine infusion in mice with cardiomyocyte-specific NCLX overexpression, which enhances _m_Ca^2+^ efflux and so limits net _m_Ca^2+^ accumulation^35^. In both studies, this increased mortality occurred prior to any overt decline in cardiac function, and we observed no additional deaths once the initial window of vulnerability had passed. We hypothesize that when _m_Ca^2+^ accumulation is attenuated and cardiomyocytes are unable to quickly adapt metabolically to meet increased energetic demands imposed by adrenergic stimulation or other stressors, the resulting energetic stress disrupts cellular ion handling and so enhances the risk for sudden cardiac death. Further supporting this model, acute (∼1-wk) transgenic cardiomyocyte MCUB overexpression to attenuate uniporter-dependent _m_Ca^2+^ uptake impairs mitochondrial metabolism and increases mortality in mice subjected to cardiac ischemia-reperfusion^29^.

Our current finding that *Mcu*-cKO mice failed to exhibit the normal physiological increase in cardiac contractility over the first few days of isoproterenol infusion (**Fig. 4 C-E**) agrees with our previous work demonstrating cardiomyocyte mtCU function to be strictly required for the heart’s acute sympathetic fight-or-flight response^19, 29^. It contrasts, though, with the proposal that slower, mtCU-independent routes of _m_Ca^2+^ uptake are sufficient to increase mitochondrial metabolism to support an increase in cardiac work rate when sympathetic stimulation persists for periods of roughly tens of minutes or longer^20^. At later timepoints, with 1-2-wks of sustained isoproterenol stimulation, we did not observe any detrimental effect of adult cardiomyocyte-specific *Mcu* deletion on cardiac function (**Fig. 4 C-E**). This finding differs from the recent report that cardiomyocyte *Mcu* deletion exaggerates cardiac dysfunction with 4-wk isoproterenol infusion^43^, though it should be noted that that study failed to directly compare appropriate αMHC-MCM controls with *Mcu*^fl/fl^ x αMHC-MCM mice, so it is difficult to distinguish the reported effects on heart function that may be specific to the loss of MCU function from effects that may be attributable to tamoxifen + Cre cardiotoxicity^45–48^. Another possible explanation for the discrepancies between our results and those of Wang et al.^43^ is the difference in timing of the experimental endpoints used. We concluded our studies after just 2-wks of isoproterenol infusion, where no decline in cardiac function was yet detected in the control αMHC-MCM genotype, whereas Wang et al. performed their measurements after 4-wks of isoproterenol treatment, a timepoint at which control animals did show appreciable cardiac decompensation.

Nevertheless, our finding that transgenic MCU overexpression clearly accelerated the progression to cardiac failure and exaggerated hypertrophy during chronic isoproterenol infusion (**Fig. 4 C-F**) argues against the notion that _m_Ca^2+^ uptake through the mtCU in cardiomyocytes strictly limits contractile dysfunction and pathological remodeling in response to chronic stress. Considering our results in light of the observations that neither global, nor cardiomyocyte-specific *Mcu* deletion, has any detrimental effect on cardiac function following a chronic increase in cardiac afterload^20, 26^, we conclude that loss of cardiomyocyte mtCU function *does not* necessarily accelerate the progression of failure in hearts subjected to chronic stress. A further point of disagreement between our model and the report by Wang et al.^43^ is that they found a significant increase in cardiac mitochondrial MCU protein expression after both 2-and 4-wks of isoproterenol stimulation, while we observed no increase in MCU protein expression following isoproterenol infusion in any genotype (**Fig. 6A, Supplemental Fig. S6A**). This calls into doubt the idea that cardiac MCU protein expression and mtCU function must increase beyond steady-state levels to enable the heart’s long-term physiological adaptation to a sustained increase in workload. Indeed, despite upregulation of cardiac MCU protein after 4-wks of isoproterenol stimulation, Wang et al^43^ also reported a profound downregulation of cardiac EMRE protein expression, which should limit _m_Ca^2+^ uptake through the mtCU, likely as a compensatory response to increased MCU expression in order to limit ongoing _m_Ca^2+^ overload.

Finally, despite our observation that mtCU function is required to increase cardiac output at the onset of adrenergic stimulation, we saw no effect of cardiomyocyte-specific *Mcu* deletion to alter the extent of hypertrophic remodeling that occurred with 14-days of isoproterenol infusion (**Fig. 4 F-G**). _m_Ca^2+^ accumulation during neurohormonal stimulation nevertheless contributes to cardiomyocyte growth and cardiac hypertrophy^35^. Thus, this result suggests that alternative, uniporter-independent modes of _m_Ca^2+^ uptake may contribute to this hypertrophic effect, even though these alternative routes of _m_Ca^2+^ loading are insufficient to support an increased cardiac work rate. Our finding that transgenic cardiomyocyte MCU overexpression exaggerated hypertrophic remodeling with isoproterenol infusion (**Fig. 4F, Fig 6E**) provides additional support for the notion that increased _m_Ca^2+^ content *per se* – regardless of the particular mechanism by which this occurs – contributes to cardiac hypertrophy.

The normal increase in cardiac contractility observed throughout the first few days of isoproterenol infusion was preserved in MCU-Tg hearts. However, within 1-2 wks of isoproterenol infusion, MCU-Tg hearts quickly decompensated towards failure (**Fig. 4, Fig. 6**). Along with a lack of early functional responsiveness to isoproterenol in *Mcu*-cKO mice, the timeframe over which detrimental effects of MCU overexpression became apparent emphasizes that continuous _m_Ca^2+^ loading shifts from being an adaptive component of the heart’s initial energetic response to stress, to eventually becoming a maladaptive response that hastens the development of contractile dysfunction. Our results regarding MCU overexpression again directly conflict with the study by Wang et al.^43^, which concluded that cardiomyocyte-specific MCU overexpression protects the heart from hypertrophy and functional decompensation with chronic 4-wk isoproterenol infusion. Differences in these studies with respect the amount of time available for compensation to occur between the induction of cardiomyocyte MCU overexpression and onset of isoproterenol administration; the dose of isoproterenol used (70mg/kg/day here, lower 10mg/kg/day in Wang et al.^43^); and the extent of MCU overexpression (only ∼4-fold in Wang et al.^43^) may help to account for these discrepancies. The greater degree of MCU overexpression achieved in our genetic mouse model, which was associated with accelerated progression towards heart failure with isoproterenol infusion, raises a critical concern for proposed therapeutic strategies that aim to boost _m_Ca^2+^ accumulation to improve cardiac energetics and redox balance in the failing heart^27, 49, 50^. *What is the safety factor for enhancing mtCU function and augmenting _m_Ca^2+^ uptake in heart disease?* Any strategy seeking to increase _m_Ca^2+^ signaling to enhance mitochondrial energetics will need to carefully balance this outcome against the risk of triggering deleterious _m_Ca^2+^ overload, especially in settings where cytosolic Ca^2+^ signaling may be chronically elevated, as occurs in numerous heart conditions. Another pertinent point that will require further study is whether enhancing mtCU activity is in fact an appropriate goal to slow the development and progression of cardiac dysfunction and/or pathological remodeling in heart disease, or if this intervention may only be appropriate for a heart that is already in fulminant or end-stage heart failure.

A key question arising from our *in vivo* results is how exactly cardiomyocyte-specific MCU overexpression accelerated contractile decompensation during isoproterenol stimulation. Genetic deletion of the mPTP regulator cyclophilin D that is required for mPTP function failed to attenuate isoproterenol-induced contractile dysfunction in MCU-Tg hearts, and surprisingly increased rather than decreased isoproterenol-induced cardiomyocyte necrosis in MCU-Tg animals (**Fig. 6**). Although deletion of cyclophilin D has been reported to sensitize hearts to physiological and pathological cardiac hypertrophy, and to exacerbate pressure overload-induced contractile dysfunction, these phenotypes were not associated with increased rates of cardiomyocyte death as measured by TUNEL staining^51^. Our current results thus raise two intriguing questions regarding the response of MCU-Tg hearts to chronic isoproterenol infusion:

1. *how does deletion of cyclophilin D exacerbate isoproterenol-induced cardiomyocyte death in MCU-Tg hearts? and 2) what is the mechanism by which chronic _m_Ca^2+^ overload caused cardiomyocyte death and contractile dysfunction, if not through classical CypD-regulated mitochondrial permeability transition?* While answering these questions is beyond the scope of the current study, several relevant points for future investigation are addressed below.

First, transient opening of the mitochondrial permeability transition pore has been proposed as an alternative physiological mechanism for Ca^2+^ to exit the mitochondrial matrix, thus mitigating the risk of deleterious cardiomyocyte _m_Ca^2+^ overload^51, 52^. Deletion of CypD alone did not cause contractile dysfunction or increase cardiomyocyte death either under basal conditions or with isoproterenol stress. Likewise, the combination of CypD deletion and transgenic MCU overexpression was not deleterious at baseline, and exacerbated cardiomyocyte death only upon chronic adrenergic stimulation (**Fig. 6**). During isoproterenol infusion, it is plausible that increased capacity for _m_Ca^2+^ uptake and diminished capacity for _m_Ca^2+^ efflux in MCU-Tg x *Ppif*^-/-^ hearts could combine to drastically augment _m_Ca^2+^ overload. Although MCU overexpression was maintained throughout 14-days of isoproterenol treatment in both MCU-Tg and MCU-Tg x *Ppif*^-/-^ hearts (**Fig. 6A, Supplemental Fig. S6A**), chronic isoproterenol infusion significantly decreased EMRE protein expression compared to sham animals in MCU-Tg, *Ppif*^-/-^ hearts. Such downregulation of EMRE is proposed as a compensatory mechanism to limit ongoing _m_Ca^2+^ overload^53^. That downregulation of EMRE with isoproterenol stimulation occurred exclusively in MCU-Tg x *Ppif*^-/-^ hearts suggests that this group experienced the greatest degree of net _m_Ca^2+^ loading, reaching _m_Ca^2+^ levels sufficient to trigger compensatory remodeling of the mtCU. We therefore hypothesize that the greater degree of cardiomyocyte death observed in MCU-Tg x *Ppif*^-/-^ hearts with chronic isoproterenol infusion is directly attributable to a greater degree of sustained _m_Ca^2+^ overload in this experimental group.

Second, the specific mechanism(s) by which exaggerated _m_Ca^2+^ overload in MCU-Tg and MCU-Tg x *Ppif*^-/-^ hearts caused cardiomyocyte death and contractile dysfunction remains to be elucidated. Since cardiomyocyte necrosis was exaggerated with CypD deletion, the mechanism for this cell death associated with chronic excess _m_Ca^2+^ loading appears not to be attributable to classical, CypD-regulated high-conductance opening of the mPTP. An alternative possibility is that chronic _m_Ca^2+^ overload in isoproterenol-treated MCU-Tg and MCU-Tg x *Ppif*^-/-^ hearts triggers mitochondrial dysfunction, permeability transition, and cell death through the activation of mitochondrial calpains, a class of calcium sensitive proteases, and an increase in local proteolysis. Such a model would predict that a greater degree of _m_Ca^2+^ loading could lead to more mitochondrial calpain activation in isoproterenol-treated MCU-Tg x *Ppif*^-/-^ mice, thus explaining the increased amount of cardiomyocyte death observed in this group. Indeed, mitochondrial calpains can elicit permeability transition in the context of acute _m_Ca^2+^ overload during cardiac ischemia-reperfusion injury^54–56^. It remains to be determined whether excessive mitochondrial calpain activity likewise contributes to mPT, cardiomyocyte death, and contractile dysfunction resulting from persistent _m_Ca^2+^ stress.

In summary, this study provides evidence for a causal role of mtCU-dependent _m_Ca^2+^ uptake in both early physiological adaptations to an increase in cardiac workload, and the maladaptive processes promoted by sustained cardiac demand that can result in heart failure. Our finding of _m_Ca^2+^-driven cardiomyocyte necrosis that is exacerbated by loss of CypD emphasizes a need for deeper understanding of alternative pathways that can trigger cardiomyocyte death and contractile dysfunction in the failing heart. It also strengthens a model in which the classical mPTP has beneficial physiological roles to limit deleterious _m_Ca^2+^ overload in the face of chronic _m_Ca^2+^ stress, distinct from its role to initiate acute cell death. Future investigations should prioritize longitudinal studies, rather than functional measurements at single endpoints, to better define the role of _m_Ca^2+^ signaling throughout the development and progression of heart disease, and to determine the appropriate stages throughout this process for therapeutic intervention with strategies targeting _m_Ca^2+^ flux. Finally, given the endogenous counterregulatory mechanisms that appear to balance the risk of _m_Ca^2+^ overload with the heart’s dependence on Ca^2+^-regulated mitochondrial energy metabolism, we propose that interventions that unidirectionally block or enhance _m_Ca^2+^ uptake in chronic heart disease may ultimately be of limited therapeutic efficacy. We instead suggest that improving the dynamic flexibility of _m_Ca^2+^ exchange by targeting the regulatory mechanisms controlling _m_Ca^2+^ flux may be a more effective strategy for the mitigation of chronic heart failure.

## AUTHOR CONTRIBUTIONS

Conception and design of research: J.F.G. and J.W.E.

Performed experiments: J.F.G., T.S.L., J.P.L., A.S.M., E.K.M., A.N.H., and P.J. Analyzed data: J.F.G., T.S.L, and J.W.E.

Interpreted results of experiments: J.F.G., T.S.L., P.J., and J.W.E. Prepared figures: J.F.G.

Drafted manuscript: J.F.G.

Edited and revised manuscript: J.F.G. and J.W.E.

Approved final version of manuscript: J.F.G., T.S.L., J.P.L., A.S.M., E.K.M., A.N.H., P.J., and J.W.E.

## Abbreviations

Ca^2+^: calcium CypD: cyclophilin D
_i_Ca^2+^: intracellular calcium
_m_Ca^2+^: mitochondrial calcium
mtCU: mitochondrial calcium uniporter channel
MCU: mitochondrial calcium uniporter
mPT: mitochondrial permeability transition
mPTP: mitochondrial permeability transition pore

