## Supplemental Appendix for "MCU gain-and loss-of-function models define the duality of mitochondrial calcium uptake in heart failure"

**SUPPLEMENTAL METHODS**

**qPCR gene expression analysis.**

cDNA was prepared and used for quantitative PCR (qPCR) as detailed previously^1^. In brief, mRNA was isolated from frozen heart tissue using a RNeasy Fibrous Tissue Kit (Qiagen #74704) and cDNA was prepared from mRNA using a High-Capacity cDNA Reverse Transcription kit (ThermoFisher #4368813), following the manufacturers’ instructions. qPCR used PowerUp SYBR Green Master Mix (Applied Biosciences #100029283) and was performed with a CFX96 Touch Real-Time PCR Detection System (Bio-Rad). Cycler conditions for qPCR were: UDG activation at 50^o^C for 2min; initial denaturation at 95^o^C for 10min; and 40 cycles consisting of 95^o^C for 15sec, 60^o^C for 30sec, and 72^o^C for 30sec. **Supplemental Table 1** includes sequences for all qPCR primers against mouse transcripts.

**Statistics.**

All data are presented as mean ± S.E.M. Statistical analyses were carried out with Prism 6.0 (GraphPad Software). A two-tailed t-test was used for direct comparisons between two groups. Welch’s correction was used in cases of unequal variance. Grouped endpoint data were analyzed by 2-way ANOVA with Sidak’s post-hoc analysis. Correlation between experimental parameters was assessed by linear regression. For all analyses, *P* values less than 0.05 were considered significant.

**Supplemental Table 1:** Sequences of qPCR primers for mouse genes.

| Gene | Forward Primer (5’🡪3’) | Reverse Primer (5’🡪3’) |
| --- | --- | --- |
| *Emre (Smdt1)* | GGGACACTCATCAGCAAGAA | CTCCCTGTGCCCTGTTAATC |
| *Mcub* | CGACAACATCGGCTTGACTA | GTGGAGCCACAGGATGAAATA |
| *Micu1* | AAGAACACTCCCTGCCATTT | GCCAGGGTCATCTGCATTAT |
| *Micu2* | GTGCTTTCTGGAGGGCTAAA | CTGCAAGTATTCCCTAAGCTATCA |
| *Micu3* | CGACCTTCAAATCCTGCCTG | TCTGCGTGCTCTGACCTTAC |
| *Mcur1* | GCTCAAAGCCCACATACTCTTA | CTACCCAGGTTAGCCCTTATTG |
| *Nclx (Slc8b1)* | GCCATCTCCACTAACCTCAAA | GGGTCTGAGAAAGCCACTAAA |
| *Letm1* | TCCTGCGTTTCCAGCTCACCAT | GTCTTCTGTGACACCGAGAGCT |
| *Rps13* | GCACCTTGAGAGGAACAGAA | GAGCACCCGCTTAGTCTTATAG |

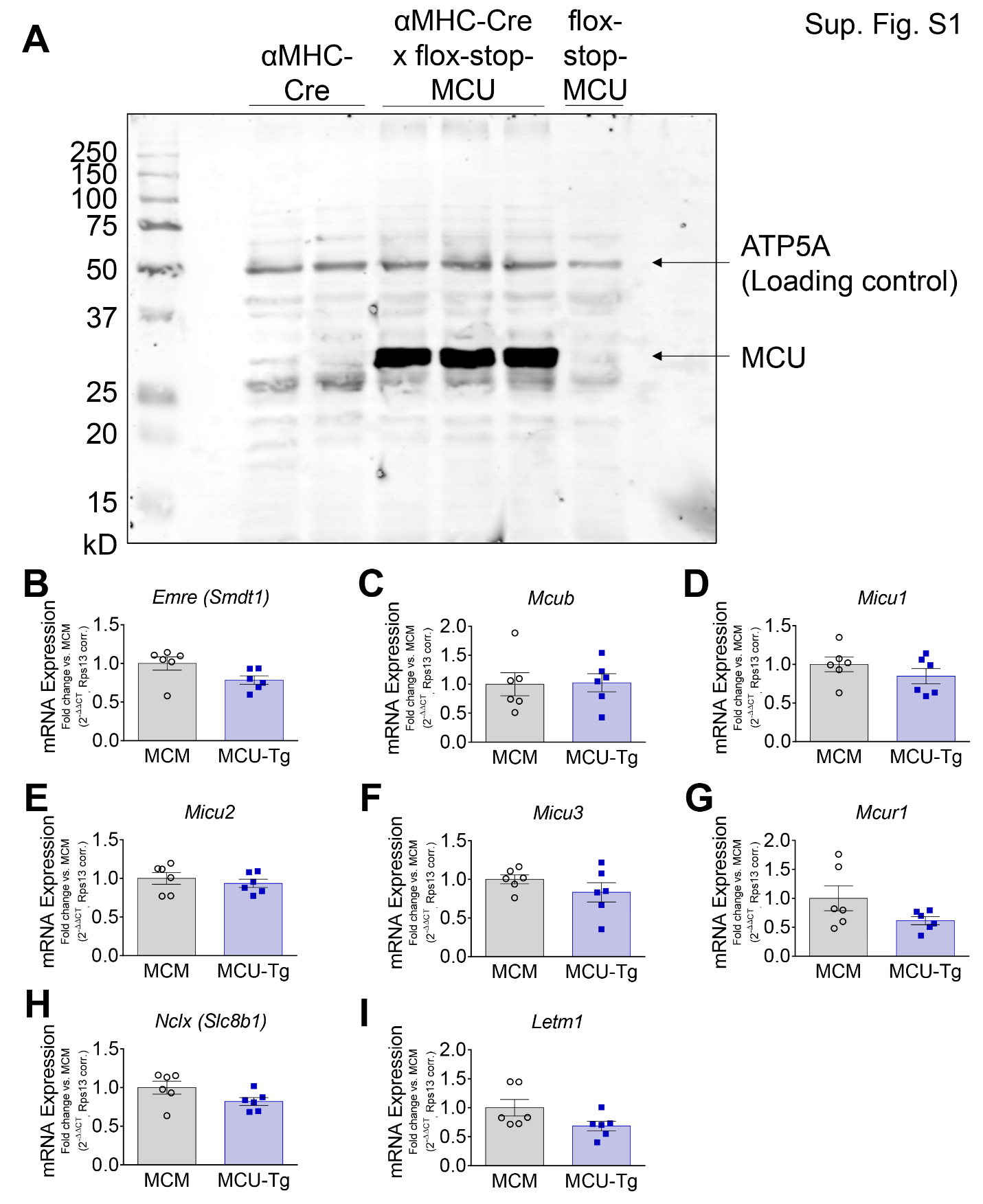

**Supplemental Figure S1: Related to Figure 1 – Mouse model of inducible, adult cardiomyocyte-specific MCU transgene expression. A)** Full-length Western blot for MCU in heart lysates from adult αMHC-Cre, αMHC-Cre x flox-stop-MCU, and flox-stop-MCU mice. The membrane was re-probed for ATP5A as a mitochondrial loading control. qPCR quantification of mRNA expression of genes encoding mitochondrial calcium uniporter complex subunits (**B**-**G**) and mitochondrial Ca^2+^ exchangers (**H**-**I**) in the hearts αMHC-MCM (MCM) and αMHC-MCM x flox-stop-MCU (MCU-Tg) mice collected 1-wk after the start of tamoxifen injections. Data analyzed by unpaired, two-tailed *t*-test. All differences between genotypes were non-significant. (*n*=6 mice/genotype).

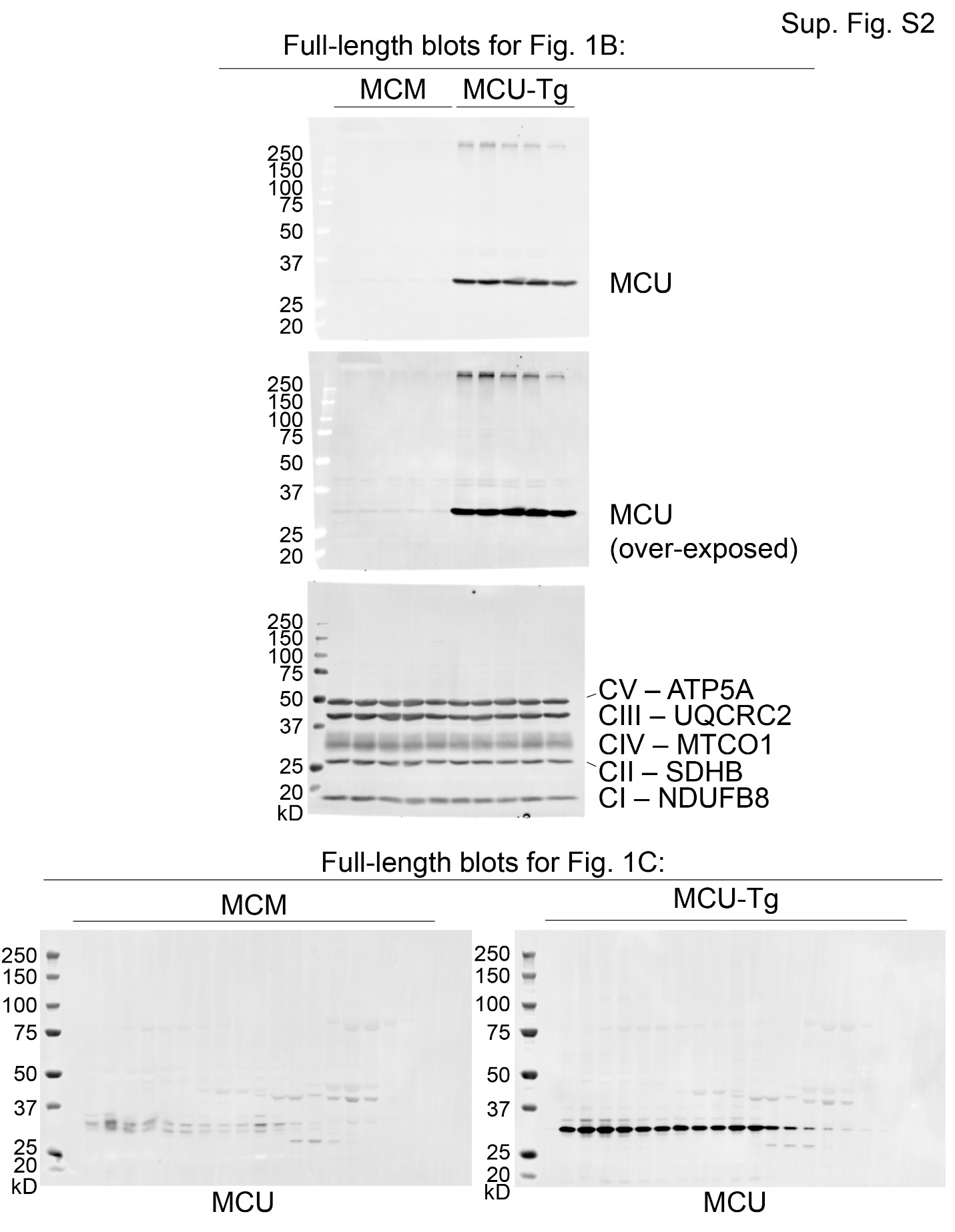

**Supplemental Figure S2:** Full-length blots for Figure 1 B and C. Two exposure times are shown for the MCU blot corresponding to Fig 1B.

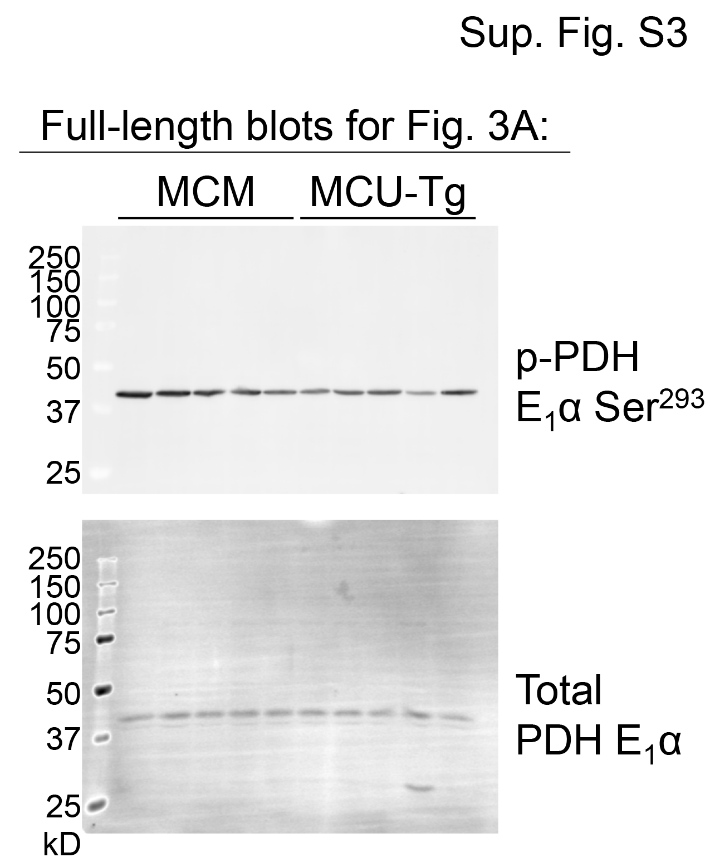

**Supplemental Figure S3:** Full-length blots for Figure 3A.

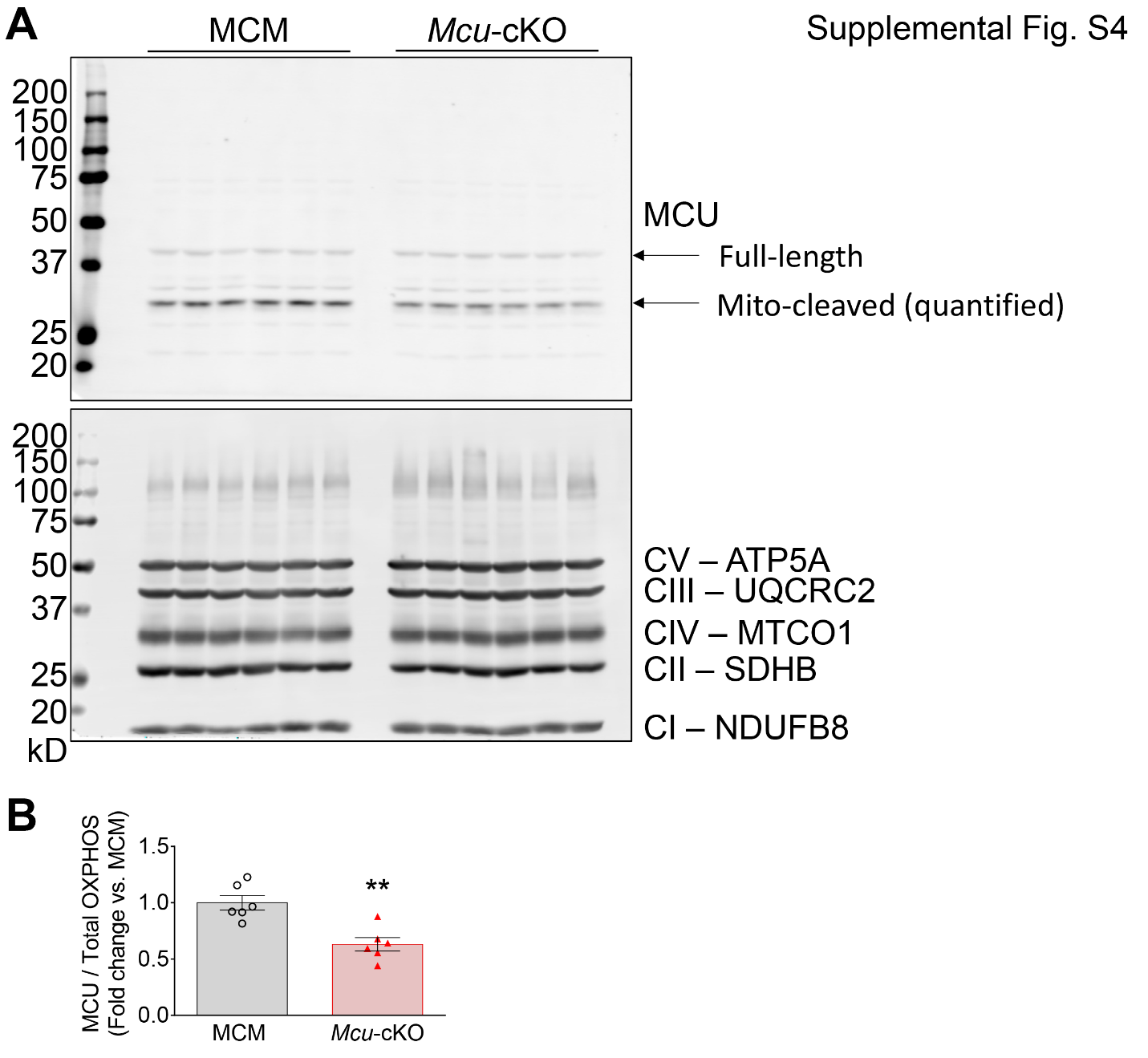

**Supplemental Figure S4: Related to Figure 4 – Cardiomyocyte *Mcu* deletion prevents isoproterenol-induced increase in cardiac contractility, while MCU overexpression predisposes to isoproterenol-induced contractile dysfunction. A**) Full-length Western blots for MCU and total OXPHOS complexes I-V protein expression in whole-heart lysates from αMHC-MCM (MCM) and αMHC-MCM x Mcu*^fl/fl^ (Mcu*-cKO) mice collected 3-wks after the start of tamoxifen. **B**) Semi-quantification of MCU protein expression normalized to total OXPHOS complexes I-V as a mitochondrial loading control. Data analyzed by unpaired, two-tailed *t*-test. ***p*<0.01. (*n*=6 mice/genotype).

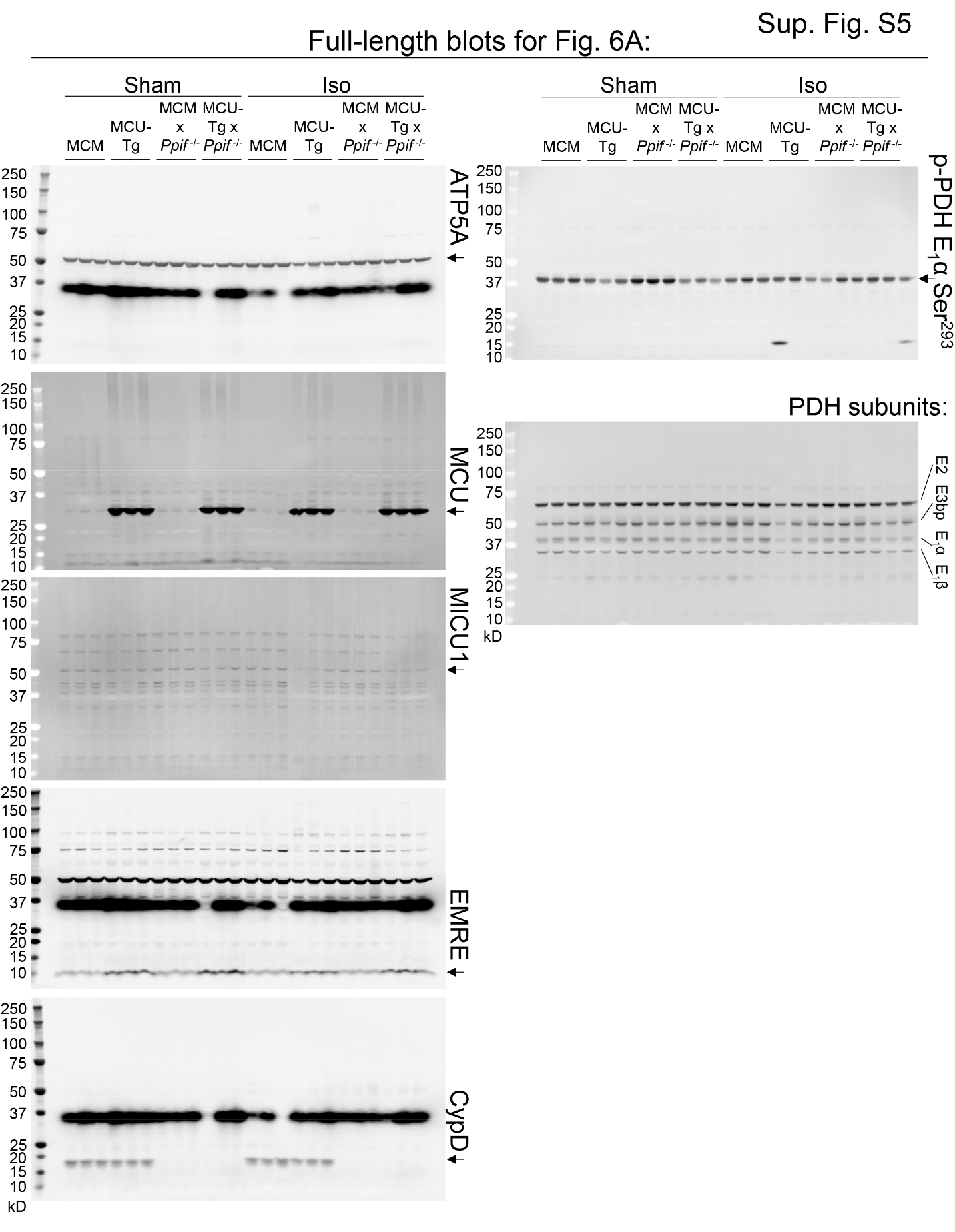

**Supplemental Figure S5:** Full-length blots for Figure 6A. Arrows indicate bands of interest.

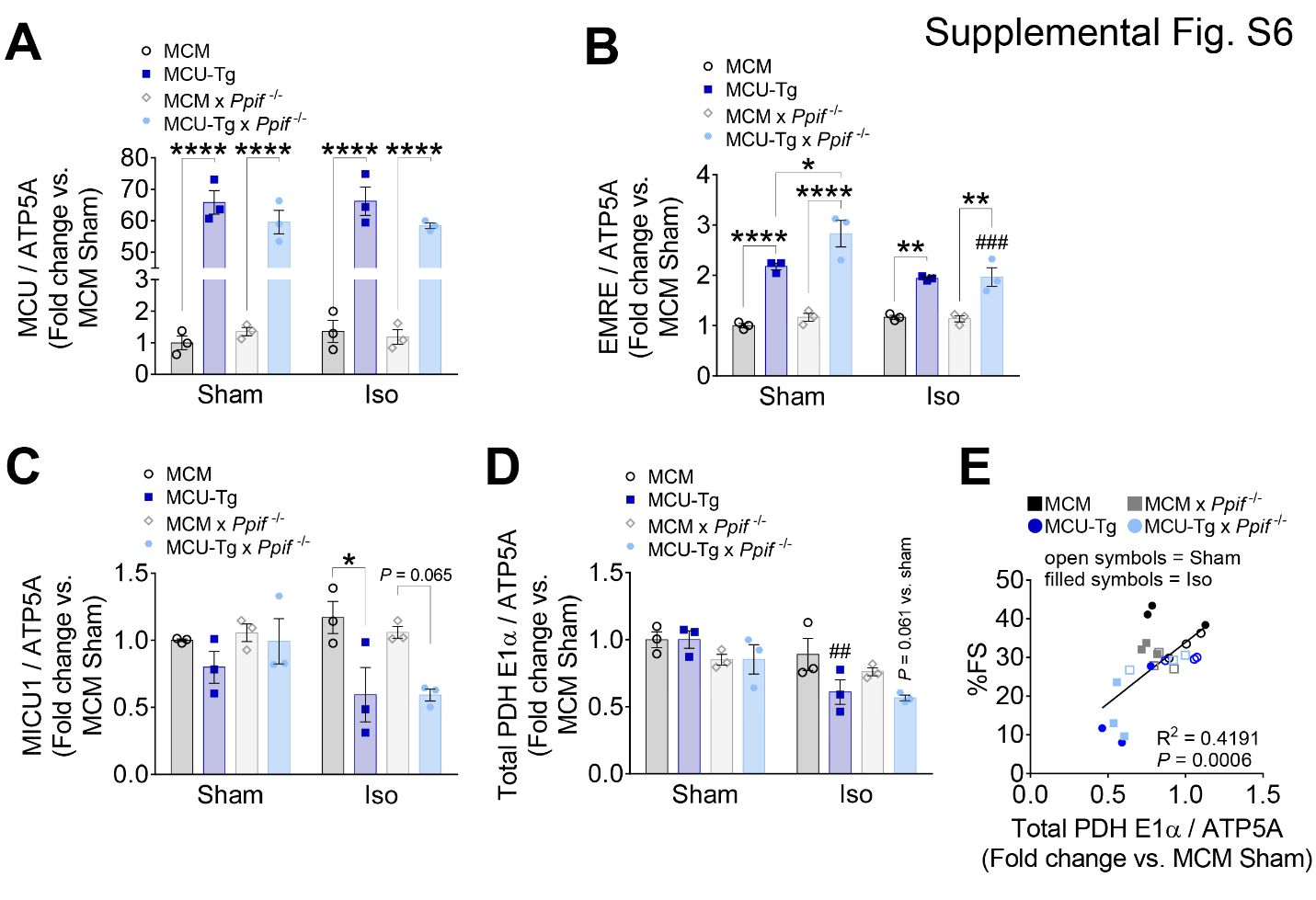

**Supplemental Figure S6: Related to Fig. 6 - Genetic inhibition of the mPTP (*Ppif*^-/-^) does not prevent contractile dysfunction, remodeling, or cardiomyocyte death in MCU-Tg hearts during chronic isoproterenol infusion.** Semi-quantification of protein expression from Fig. 6A for MCU (**A**), EMRE (**B**), MICU1 (**C**), and total PDH E_1_α subunit (**D**), each normalized to ATP5A as a mitochondrial loading control. Data analyzed by 2-way ANOVA with Sidak’s post-hoc test. **p*<0.05, ***p*<0.01, *****p*<0.0001 between genotypes; ^##^*p*<0.01, ^###^*p*<0.001 vs. Sham. (*n*=3 mice/group). **E**) Linear regression analysis of left ventricular fractional shortening (%FS) vs. total PDH E_1_α subunit expression after 14-day isoproterenol infusion. (*p* = 0.0006; *n*=3 mice/group; 24 mice total).
